# A novel stem cell type at the basal side of the subventricular zone maintains adult neurogenesis

**DOI:** 10.1101/2020.11.20.391102

**Authors:** Katja Baur, Yomn Abdullah, Claudia Mandl, Gabriele Hölzl-Wenig, Yan Shi, Udo Edelkraut, Priti Khatri, Anna M. Hagenston, Martin Irmler, Johannes Beckers, Francesca Ciccolini

## Abstract

According to the current consensus, neural stem cells (NSCs) apically contacting the lateral ventricle generate differentiated progenitors by rare asymmetric divisions or by relocating to the basal side of the ventricular-subventricular zone V-SVZ. Both processes will then ultimately lead to the generation of adult-born olfactory bulb (OB) interneurons. In contrast to this view, we here found that adult-born OB interneurons largely derive from an additional NSC type resident in the basal V-SVZ. Despite being both capable of self-renewal and long-term quiescence, apical and basal NSCs differ in Nestin expression, primary cilia extension and frequency of cell division. The expression of Notch-related genes also differed between the two NSC groups and Notch-activation was greatest in apical NSCs. Apical downregulation of Notch-effector Hes1 decreased Notch activation while increasing proliferation across the niche and neurogenesis from apical NSCs. Underscoring their different roles in neurogenesis, lactation-dependent increase in neurogenesis was paralleled by extra activation of basal but not apical NSCs. Thus, basal NSCs support OB neurogenesis whereas apical NSCs impart Notch-mediated lateral inhibition across the V-SVZ.

## Introduction

The adult mammalian brain, despite being generally unable to repair and regenerate, encompasses self-renewing and multipotent neural stem cells (NSCs) (Gage, 2002; Obernier & Alvarez-Buylla, 2019). In mice NSCs were first retrospectively identified in clonal assays, demonstrating that epidermal growth factor (EGF) elicits the proliferation of rare self-renewing and multipotent adult brain cells (Reynolds & Weiss, 1992). Thereafter, it was shown that NSCs residing in the subgranular zone (SGZ) of the hippocampus and in the ventricular-subventricular zone (V-SVZ) lining the lateral ventricles sustain neurogenesis throughout adulthood (Ming & Song, 2011). The largest neurogenic niche in the adult murine brain is the V-SVZ, where adult-born neuroblasts migrate along the rostral migratory stream to the olfactory bulb (OB) where they differentiate into GABAergic and, to a lesser extent, glutamatergic interneurons(Brill *et al*, 2009; Lois *et al*, 1996; Merkle *et al*, 2007). Underscoring its functional relevance, adult OB neurogenesis contributes to odor recognition (Feierstein, 2012) and is modulated by conditions engaging the olfactory function like the behaviour associated with mating and parenting (Furuta & Bridges, 2005; Shingo *et al*, 2003).

Most cells in the adult murine brain originate from radial glia progenitors (Anthony *et al*, 2004; Malatesta *et al*, 2003). Radial glia cells displaying apical-basal polarity are first generated at the onset of neurogenesis between embryonic day (E) 10 and 12 (Kriegstein & Gotz, 2003). They are characterized by an apically located centrosome as well as Nestin (Hockfield & McKay, 1985) and Prominin-1 (Weigmann *et al*, 1997) expession. The astroglial marker glial fibrillary acidic protein (GFAP) is first expressed in this population after birth, when many radial glia cells transform into astrocytes(Alves *et al*, 2002; Kalman & Pritz, 2001; Misson *et al*, 1991).

Radial glia cells also give rise to adult NSCs (Merkle *et al*., 2007; Ortiz-Alvarez *et al*, 2019), which maintain apical-basal polarity as well as Nestin, GFAP and Prominin-1 expression (Mirzadeh *et al*, 2008; Shen *et al*, 2008; Tavazoie *et al*, 2008). In the adult V-SVZ, proliferating NSCs and more differentiated intermediate progenitors, including transit amplifying progenitors (TAPs) and pre-neuroblasts, display high levels of EGF receptor (EGFR) which is downregulated during neuronal differentiation (Carrillo-Garcia *et al*, 2010; Cesetti *et al*, 2009; Codega *et al*, 2014; Doetsch *et al*, 1999; Khatri *et al*, 2014).

Although genetic tagging has shown that cells displaying GFAP promoter activity contribute to neurogenesis in the olfactory bulb (OB) (Beckervordersandforth *et al*, 2010; Doetsch *et al*., 1999; Garcia *et al*, 2004; Weber *et al*, 2011), it is unclear whether these tagged cells represent apical radial glia-like NSCs (Joppe *et al*, 2020). Indeed, during embryonic development, apical radial glia can give rise to basal radial glia that lack apical attachment, undergo mitosis at the basal side of the niche and are critical for the numeric expansion of neurons in gyrencephalic brains (Florio & Huttner, 2014; Penisson *et al*, 2019). The generation of basal progenitors from apical NSCs has also been observed in the V-SVZ of neonatal(Alves *et al*., 2002; Tramontin *et al*, 2003) and in the older mice (Obernier *et al*, 2018). However, in both cases, rather than asymmetric self-renewing division of apical progenitors, basal progenitor generation involves apical detachment and migration to the basal V-SVZ, suggesting that neurogenesis in the postnatal V-SVZ is associated with NSC consumption.

In contrast to this model, we here show the presence of NSC lacking apical attachment which and capable of undergoing long-term quiescence, activation, and self-renewal at the basal side of the postnatal V-SVZ. From birth onwards, these basal NSCs are the largest NSC pool in the postnatal V-SVZ and the main source of adult-born OB interneurons. Moreover, the temporary increase in proliferation, which occurs in the V-SVZ following lactation (Shingo *et al*., 2003), is sustained by activation of basal NSCs and does not affect the number of total apical NSCs. Our data show that rather than directly contributing to neurogenesis, apical NSCs contribute to niche homeostasis by regulating levels of Notch-mediated lateral inhibition across the V-SVZ.

## Results

### The majority of hGFAP-tagged progenitors display no apical membrane and no Prominin-1-immunoreactivity

Adult NSCs are identified based on the expression of markers and functional characteristics. Therefore, to directly investigate NSCs we first took advantage of a genetic mouse model enabling *in vivo* tagging of NSCs. Adult NSCs were analysed in 8-weeks-old (8W) hGFAP;H2B-GFP mice, in which the tetracycline-responsive transactivator (tTA) expressed from a human GFAP promoter controls in a Tet-off fashion the transcription of histone 2B fused to the green fluorescent protein (H2B-GFP)(Luque-Molina *et al*, 2017). Cells contacting the lateral ventricle with the apical membrane were labelled by applying DiI on the apical side of the V-SVZ before tissue dissection (Fig. 1A, B) and additional Prominin-1 immunostaining before FACS analysis (Fig. 1D-F), as previously reported (Khatri *et al*., 2014). As expected, H2B-GFP expressing (G^+^) NSCs represented a small minority of the total viable cells (Fig. 1A). Nevertheless, compared to the remaining G^-^ cells, they were significantly enriched in DiI^+^ apical cells (Fig. 1B), representing 23.54% ± 2.62 of the total DiI^+^ cells (Fig. 1C). Similarly, the incidence of Prominin-1-immunopositive (P^+^) cells was higher in the G^+^ population than in the remaining G^-^ cells (Fig. 1D, E), and G^+^P^+^ cells were composed mostly of DiI^+^ cells (Fig. 1F), which is consistent with Prominin-1 being prominently expressed in apical V-SVZ cells. However, the majority of H2B-GFP tagged cells neither expressed Prominin-1 (Fig. 1E) nor displayed DiI labelling (Fig. 1B), indicating that they are basal progenitors lacking an apical attachment.

**Figure 1:**
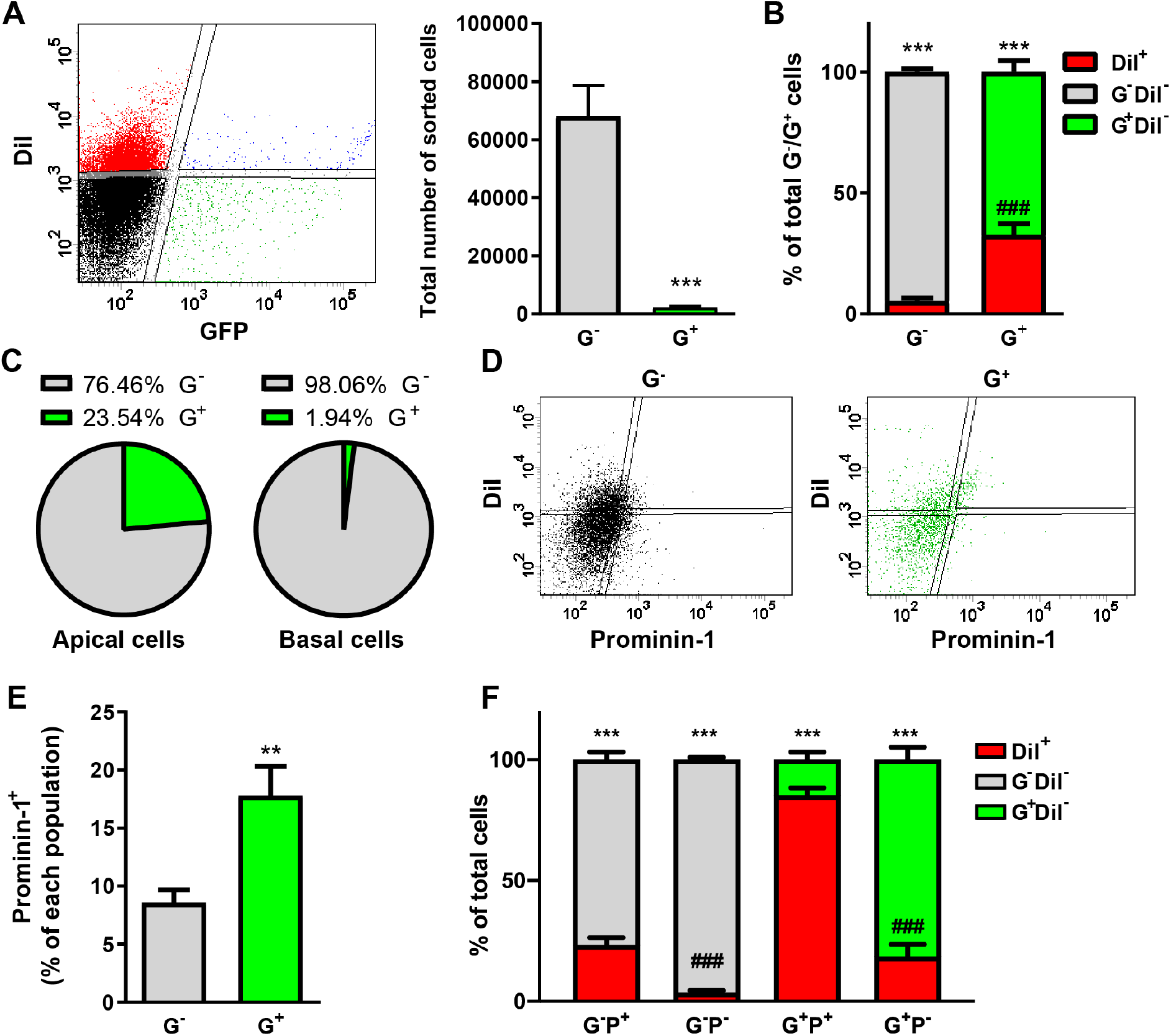
Most hGFAP-tagged cells display neither an apical membrane nor Prominin-1 expression. (A) Representative FACS plot illustrating the distribution of dissociated V-SVZ cells according to their H2B-GFP (GFP) and DiI fluorescence (left panel). Quantitative analysis in the right panel shows the total number of V-SVZ cells expressing (G^+^) or not expressing H2B-GFP (G^-^). (B) Quantitative analysis of the percentage of DiI-labeled (DiI^+^) apical cells within G^-^ and G^+^ populations (C) and of the percentage of G^-^ and G^+^ amongst total apical (DiI^+^) and basal (DiI^-^) V-SVZ cells. (D) Representative FACS plot of G^-^ cells (left panel) and G^+^ cells (right panel) illustrating their distribution according to Prominin-1 and DiI fluorescence. (E) Quantification of the percentage of Prominin-1 immunopositive cells in G^-^ and G^+^ populations. (F) Quantitative analysis of the percentage of DiI^+^ apical cells in each G^-^ and G^+^ population sorted additionally on the basis of Prominin-1 (P) expression. Data is represented as mean ± SEM. N=29. * indicates significance between apical (DiI^+^) and basal (DiI^-^) cells within the same cell population and # indicates significance between G^-^ and G^+^ in (B) and between G^-^P^+^ and G^-^P^-^ or G^+^P^+^ and G^+^P^-^ in (F). **p<0.01, ***/ ### p<0.001.

Label-retention is a characteristic of adult stem cells. To enrich for label-retaining cells within the reporter expressing population, we next investigated the expression of NSC markers within G^+^ cells in untreated mice and in mice that had been administered doxycycline for 30 days before analysis (Fig. 2A, B). Labelled nuclei were considered to be apical progenitors if they were localized at a maximal distance from the ventricle of 10 μm along the apical-basal axis. This criterion was established upon analyses of the position of apical cells tagged by adeno associated virus (AAV) delivered within the ventricular space (supplementary Fig. S1) (Luque-Molina *et al*, 2019). Besides GFAP, we also investigated the expression of Nestin (Fig. 2A, B), which is particularly observed in actively cycling NSCs (Codega *et al*., 2014) and of the Sex-determining region Y (SRY)-box transcription factor 9 (Sox9), a marker of multipotent NSCs (Scott *et al*, 2010) (supplementary Fig. S2). Consistent with our FACS data showing a lack of apical characteristics in most tagged cells, in the absence of doxycycline, most G^+^ cells were localized at the basal side of the niche (Fig. 2B). At this side, only few Nestin immunoreactive G^+^ cells were observed, especially upon doxycycline administration. Similarly, a higher percentage of apical G^+^ cells were immunopositive for GFAP and Sox9 than were basal G^+^ cells, but these differences were not significant (Fig. S2A-C). Finally, both groups of G^+^ cells expressed similar transcripts level of the Leucine-rich repeats and immunoglobulin-like domains 1 (Lrig1) gene (Fig. SD), which has been recently established as a marker for the prospective identification of NSCs in the V-SVZ (Nam & Capecchi, 2020).

**Figure 2:**
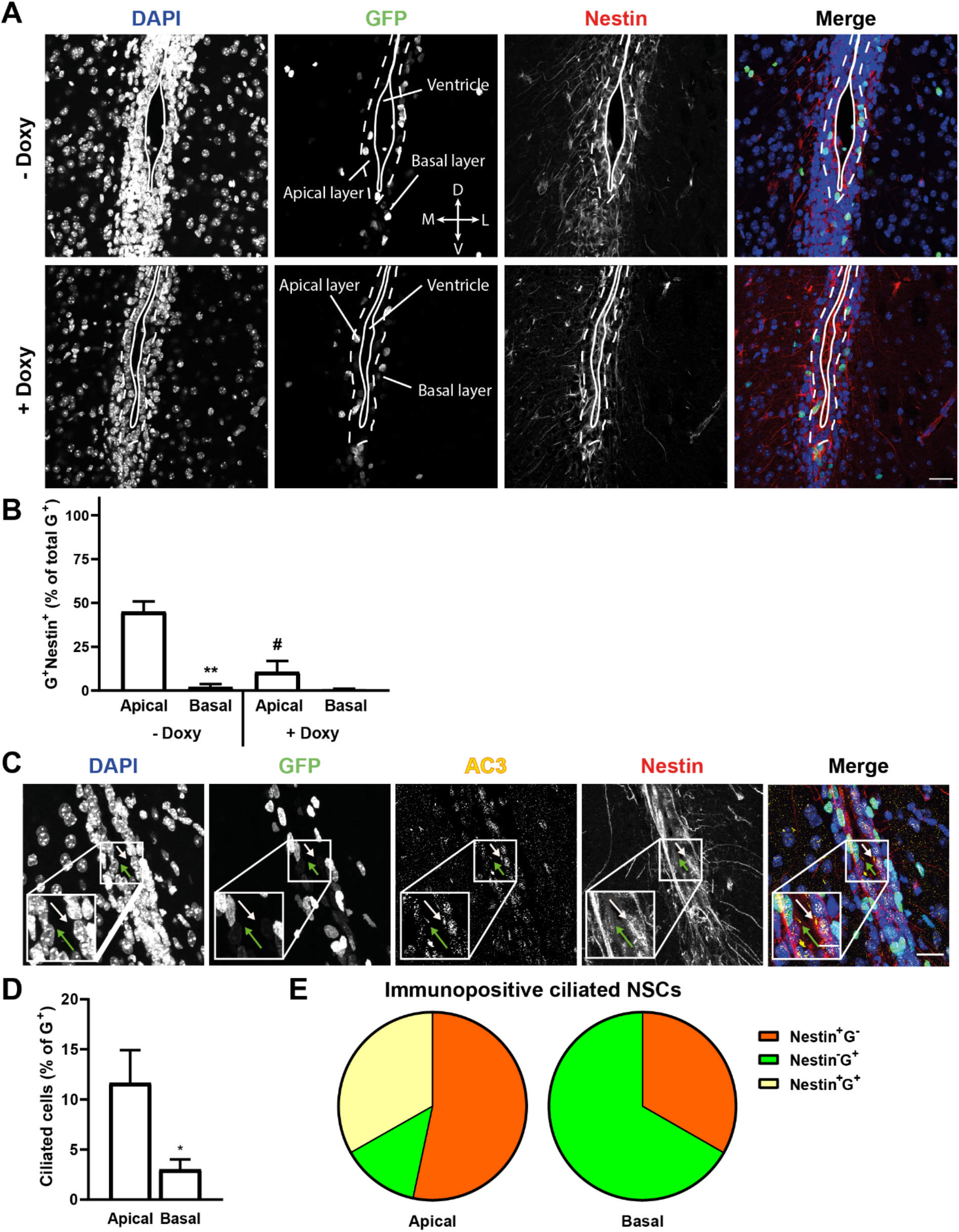
Basal hGFAP-tagged cells in the adult SVZ do not express Nestin and rarely extend primary cilia. (A) Representative confocal photomicrographs showing representative examples of coronal slices of the V-SVZ obtained from 8-weeks-old hGFAP;H2B-GFP mice treated with and without doxycycline (doxy) for 30 days as indicated. H2B-GFP (GFP) expression is shown in green, Nestin immunostaining in red and DAPI counterstaining of the nuclei in blue. Crossing arrows indicate slice directions: M = medial, D = dorsal, V = ventral, L = lateral. Continuous and dotted lines show the region where the nuclei of apical cells were counted. (B) Quantification of the proportions of Nestin immunopositive (Nestin^+^; yellow bars) and immunonegative (Nestin^-^; green bars) apical and basal G^+^ cells in doxycycline-treated and untreated animals. N=3 (no Doxy), N=4 (Doxy) (C) Representative confocal photomicrographs of coronal slices of hGFAP;H2B-GFP mice not treated with doxycycline and immunostained to illustrate the expression of H2B-GFP (GFP; green), Nestin (red) and AC3 (yellow) immunoreactivity and DAPI (blue) counterstaining of the nuclei. Arrows indicate Nestin-positive and GFP-positive cells with a primary cilium. (D) Quantification illustrating the percentage of ciliated (AC3^+^) apical and basal G^+^ cells. (E) Pie charts represent antigenic characteristics of apical and basal ciliated cells. N=7. Scale bar = 30 μm; 10 μm for insert in (C). Bars represent mean ± SEM. * and # indicate significance: between apical and basal (*) and between doxycycline-treated and nontreated counterpart (#); */# p<0.05, **p<0.01.

Primary cilia are closely linked to cell cycle progression and signal transduction (Goto *et al*, 2013). Since not all adult quiescent NSCs in the V-SVZ display primary cilia (Khatri *et al*., 2014), we next used adenylate cyclase 3 (AC3) as a primary cilia marker (Bishop *et al*, 2007; Monaco *et al*, 2019) in combination with Nestin and H2B-GFP to investigate presence of primary cilia in apical and basal progenitors (Fig. 2C, E). Ciliated progenitors represented a small subset of G^+^ tagged progenitors in the V-SVZ (Fig. 2D) especially at the basal side of the niche. However, at either side of the niche, G^+^ cells still represented the majority of ciliated progenitors with variation in percentages and Nestin expression (Fig. 2C-E). A similar association between apical NSCs and primary cilia was observed also in WT mice, where ciliated cells were enriched within DiI^+^ labelled apical cells (supplementary Fig. S3), and it is consistent previous findings (Beckervordersandforth *et al*., 2010). Notably in the latter study, using a similar approach to identify NSCs, it was proposed that hGFAP-tagged cells devoid of Prominin-1 immunoreactivity in the V-SVZ represent niche astrocytes. This conclusion was supported by gene expression analysis, showing that Prominin^+^/GFP^+^ cells display a gene expression profile different from that of GFP^+^ astrocytes isolated from the Diencephalon. However, the expression profile of GFP^+^ cells isolated from the V-SVZ was not characterized in detail in this previous study. We therefore here analyzed again these previously published data to directly compare the profiles of Prominin^+^/GFP^+^ and GFP^+^ cells isolated from the V-SVZ. A principal component analysis (PCA) of these populations showed that their transcriptomes are similarly divergent from those of diencephalic astrocytes (supplementary Fig. S3C). Moreover, analysis of differentially regulated genes highlighted significant regulation of genes associated with chromosome replication and mitosis (Supplementary table 1 and 2). The signalling pathways activated in Prominin^-^/GFP^+^ cells also included those involving the epidermal growth factor (EGF) and the anti-angiogenic pigment epithelium-derived factor (PEDF), supplementary tables 1, 2, previously shown to act as a niche-derived growth factor promoting NSC self-renewal (Ramirez-Castillejo *et al*, 2006). Taken together, these data indicate that Prominin-1 immunonegative h-GFAP tagged cells may also contain NSCs.

### Both apical and basal hGFAP-tagged progenitors display hallmarks of NSCs

To further characterize the stemness of the basal G^+^ cells, we next investigated whether basal progenitors undergo quiescence. For this analysis, we measured label-retention after 30 day-doxycycline treatment and Ki67 expression in each progenitor cell population. Since the fluorescent nuclear signal disappears after five cell divisions (Waghmare *et al*, 2008), a relative decrease in the number of total apical or basal G^+^ cells upon doxycycline administration reflects the relative number of proliferating cells at the start of doxycycline treatment. In the same vein, the lack of a decrease in the number of G^+^ cells would indicate cellular quiescence. Treatment with doxycycline led to a small reduction in the number of G^+^ cells, which was significant only within the subset of basal G^+^ progenitors (Fig. 3A, B). This shows that most basal and especially apical progenitors are slow cycling label-retaining cells. In line with this idea, the majority of apical and basal G^+^ cells did not display Ki67 immunoreactivity (Fig. 3C), confirming that in both groups the majority of the cells are non-cycling quiescent progenitors. Notably, apical and basal G^+^ progenitors contained a similar subset of Ki67^+^ cycling cells, indicating that, rather than reflecting differences in the proportion of quiescent cells, the lower number of label-retaining cells in basal than in apical G^+^ progenitors could be an indication of faster cell cycle kinetics in basal G^+^ progenitors. Consistent with this hypothesis, Ki67 expression was significantly reduced in basal but not apical G^+^ cells upon doxycycline administration, an observation that would be expected if basal Ki67^+^ cycling cells underwent more rounds of cell divisions during the chasing period.

**Figure 3:**
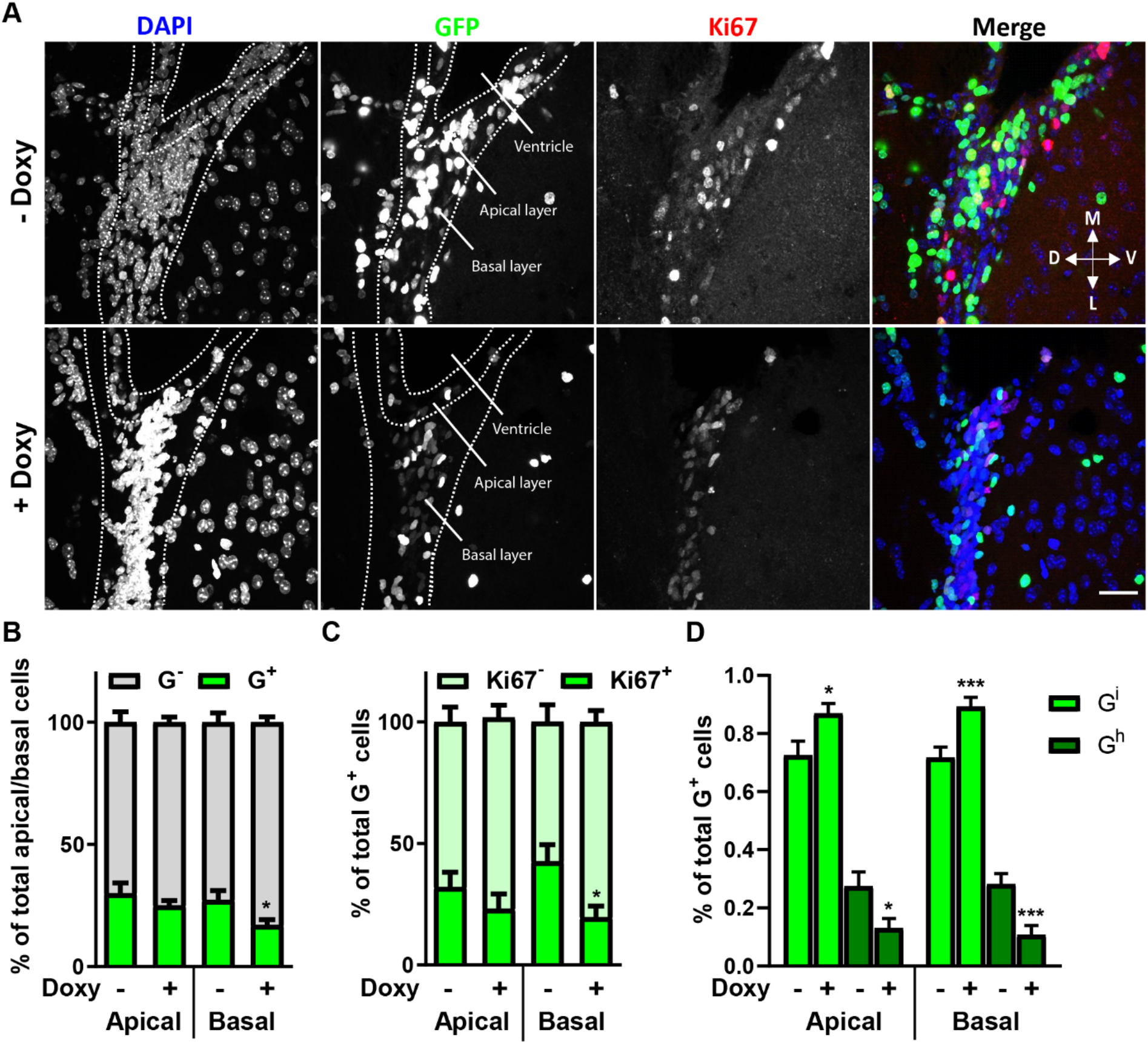
Both apical and basal hGFAP-tagged cells display NSC characteristics. (A) Repesentative confocal photomicrographs of coronal sections of the V-SVZ of 8W hGFAP;H2BGFP mice, which had been administered doxycycline (Doxy) as indicated. Nuclear H2B-GFP (GFP) expression and Ki67 immunoreactivity are shown in green and red, respectively. DAPI counterstaining of the nuclei is blue. Scale bar = 30 μm. (B-D) Quantitative analyses of apical and basal cells expressing H2B-GFP (G^+^) or not (G^-^) (B), of Ki67 immunoreactivity in apical and basal G^+^ cells (C), and of apical and basal cells expressing intermediate (G^i^) or high (G^h^) GFP levels with and without doxycycline (Doxy) treatment (D). Data is represented as mean ± SEM N≥4. * indicates significance between doxycycline-treated and -non-treated specimens within apical or basal cell populations. * p<0.05, ***p<0.001.

Self-renewal is a defining property of stem cells and we next investigated self-renewal of apical and basal G^+^ cells *in vivo* using hGFAP;H2B-GFP mice. We reasoned that in the absence of doxycycline, the expression of the H2B-GFP reporter would be downregulated only in cells undergoing non-renewing cell divisions. Thus, in control untreated mice, self-renewal would maintain high levels of fluorescence expression in the pool of G^+^ cells, whereas non-renewing cell division, comparable to doxycycline administration, would cause a gradual dilution in fluorescence levels at each cell division. Therefore, to measure the self-renewal rate, we compared the percentage of G^+^ cells expressing high (G^h^) or intermediate (G^i^) fluorescence levels, as described before(Luque-Molina *et al*., 2017), see also “Methods” section, in sorted apical P^+^G^+^ and basal P^-^G^+^ progenitors (Fig. 3D). Compared to doxycycline-treated animals, control untreated mice displayed a consistent increase in the percentage of G^h^ cells within apical and especially basal progenitors, indicating that both cell groups undergo self-renewing divisions *in vivo*.

Since neurosphere formation is a hallmark property of proliferating adult NSCs (Codega *et al*., 2014; Khatri *et al*., 2014), we next tested the ability of apical and basal neural G^+^ precursors to form clonal neurospheres (supplementary Fig. S4). For this analysis, cells were additionally sorted between those expressing high (E^h^) and low (E^l^) EGFR levels, as the first are enriched in clone forming cells compared to the latter(Ciccolini *et al*, 2005). In the absence of doxycycline, clones were generated by both basal G^+^P^-^E^h^ and G^+^P^-^E^l^ progenitors, albeit with different frequency. Instead at the apical side, consistent with previous observations in WT precursors (Khatri *et al*., 2014), only G^+^P^+^E^h^ progenitors underwent clone formation, indicating that this population maintains a slow rate of cell division also *in vitro*. Moreover, when G^+^ cells were isolated from mice which had been treated with doxycycline for one month, clone formation was drastically reduced and almost exclusively observed in the group of G^+^P^-^ E^h^ cells, in line with the notion that quiescent NSCs are not capable of forming clones and that most activated NSCs display high levels of EGFR. Taken together, these data show that although basal G^+^ progenitors undergo faster cell cycle kinetics than apical G^+^ cells, they are not rapidly proliferating intermediate precursors. On the contrary, our data show that basal G^+^ precursors display characteristics of adult NSCs, including the ability to undergo quiescence, self-renewal, and clone formation when they are in an active state. Therefore, they will be hereafter referred to as basal NSCs.

### Basal NSCs represent the predominant NSC population throughout postnatal life

It has been recently shown that with age, apical NSCs detach from the ventricular surface, giving rise to basal progenitors (Obernier *et al*., 2018). Therefore, we next investigated if basal NSCs are generated from apical NSCs during adulthood or if they are already present early on in the neonatal brain. We first analysed the presence of apical and basal NSCs in slices of the V-SVZ of 1W-old hGFAP;H2B-GFP mice (supplementary Fig. S5). Quantitative analysis showed not only the presence of both apical and basal G^+^ NSCs already at this early age, but also characteristics between the two groups with respect to Nestin expression (supplementary Fig. S5A, B), ciliation (supplementary Fig. S5A-C) and prevalence of G+ cells (supplementary Fig. S5D) similar in trend to those observed in adult apical and basal adult populations. However, independent of the niche location, apical neonatal cells displayed a greater proportion of G^+^ cells co-expressing Nestin and extending a primary cilium than the 8W-old counterpart. Taken together, these data show that cells with defining characteristics of basal and apical NSCs are present in neonatal mice.

We next investigated age-related changes in the number of apical and basal NSCs and their propensity to undergo activation. To this end, we quantified the number of apical G^+^P^+^ and basal G^+^P^-^ progenitors (Fig. 4A left panel) and the proportion of each population expressing high EGFR levels at the cell surface (E^h^; Fig. 4A right panel) at various postnatal ages. Consistent with previous observations in WT mice (Carrillo-Garcia *et al*., 2010), the number of basal and apical NSCs decreased with age (Fig. 4A left panel). However, for both apical and basal NSC, the sharpest and only significant drop in NSC numbers occurred between 1 and 2 weeks of postnatal age, similar to what was recently reported for hippocampal NSCs(Harris *et al*, 2021). In contrast, a significant decrease in the proportion of activated NSCs was observed first in 8W animals, affecting basal NSCs only, and thereafter in 25W mice for both apical and basal NSCs (Fig. 4A). We next determined the effect of age on the potential of proliferation and self-renewal of apical and basal NSCs with clonal assays (Fig. 4B-D). For this analysis, apical and basal NSCs were sorted from the V-SVZ of 8W and 25-30W old animals by flow cytometry, based also on DiI labelling of the apical membrane as well as EGFR and Prominin-1 expression. Consistent with our observations above, age led to a decrease in the number of activated G^+^P^-^E^h^ cells within the basal (D^-^) but not apical (D^+^) pool of NSCs (Fig. 4B). At both ages, all types of NSCs generated clones and in line with our findings *in vivo*, they were also able to undergo self-renewal *in vitro*, determined by the number of green cells per clone (Fig. 4C and D). However, unlike the remaining populations, apical G^+^P^+^E^h^ NSCs underwent very limited selfrenewal (Fig. 4C) and generated only small clones when isolated from older mice (Fig. 4D). A similar trend towards a decrease in the ability of generating large clones was also observed for basal G^+^P^+^E^h^ NSCs. Moreover, in both apical and basal NSCs age led to a significant increase in the proportion of G^+^P^-^E^h^ cells capable of generating large clones (Fig. 4D), suggesting that with age, more proliferating precursors loose Prominin-1 expression at the cell surface. Taken together, these data show that the basal NSCs represent the major NSC population in the postnatal V-SVZ throughout adulthood and that they are endowed with greater self-renewal potential than their apical counterpart. Within the time-frame analysed, a strong reduction in the number of basal and especially apical NSCs was only observed after the first postnatal week. Thus, rather than NSC numbers or self-renewal potential, age affects the ability to express high levels of EGFR in both groups of NSCs and the proliferation potential uniquely of apical NSCs.

**Figure 4:**
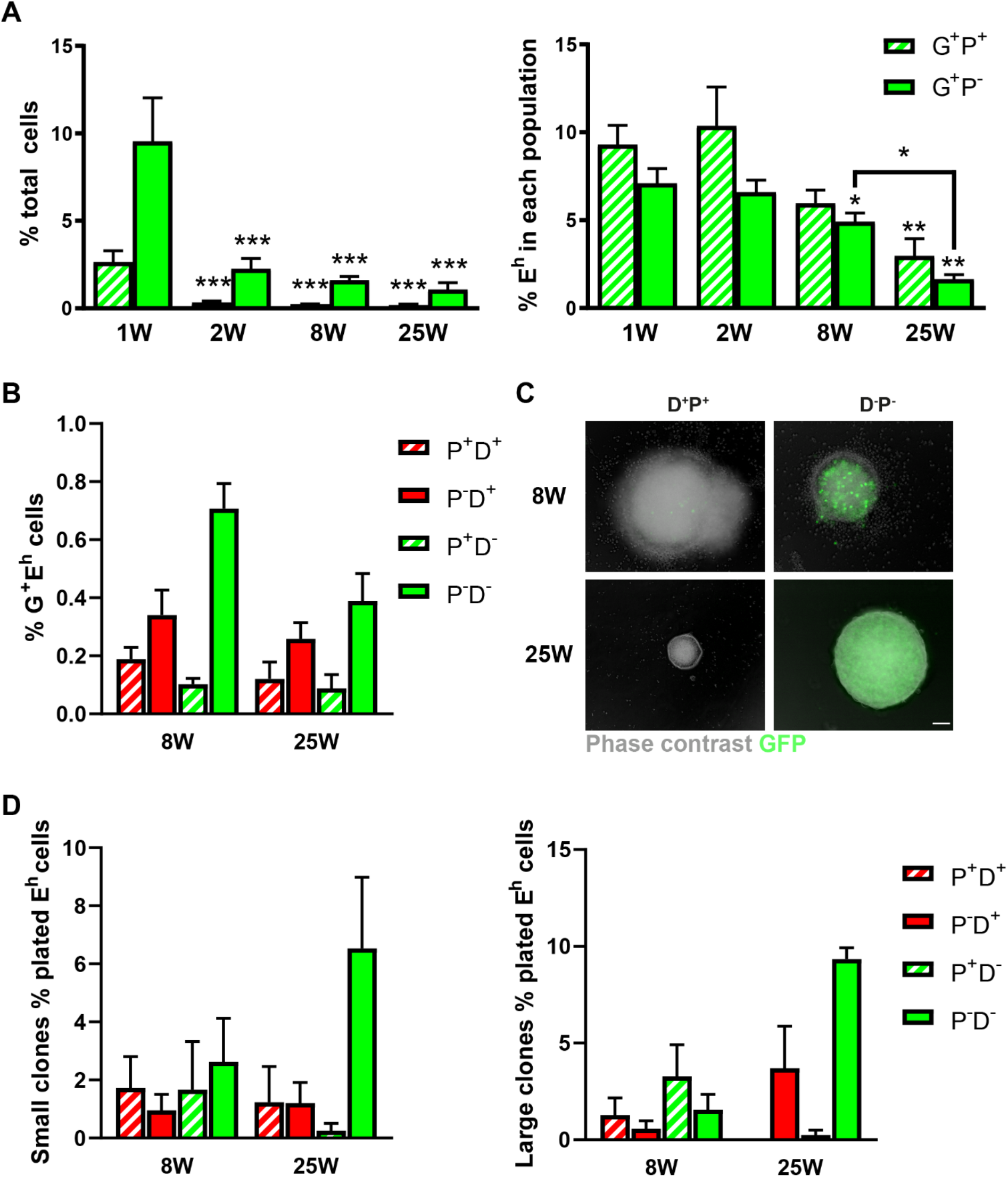
Basal NSCs represent the predominant stem cell population in the adult mouse brain. (A) Quantification of the flow cytometry analysis of the number of apical Prominin-1 immunopositive (G^+^P^+^) and basal Prominin-1 immunonegative (G^+^P^-^) H2B-GFP-expressing cells (left panel) and of the percentage of activated cells expressing high levels of EGFR (E^h^) in each population (right panel) at the various ages as indicated. (B) Quantitative analysis of the percentage of DiI-labelled apical (D^+^) and unllabelled basal (D^-^) G^+^ E^h^ cells expressing Prominin-1 (P^+^) or not (P^-^) in the V-SVZ of 8- and 25-weeks-old animals. (C) Representative micrographs of clonal neurospheres generated from G^+^E^h^ cells isolated at the indicated ages and displaying H2B-GFP (GFP), DiI labelling and Prominin-1 immunoreactivity. Scale bar = 100 μm. (D) Quantification of the percentage of the indicated subsets of G^+^E^h^ cells generating small (left panel) and large (right panel) clones after sorting from 8W or 25W animals. Bars represent mean ± SEM. N.D. = none detected. N≥4. * indicates in (A) significant difference from the corresponding 1W population (on top of bars) or between ages (lines) as indicated, in (B) and (D) from the respective population at 8 W. * p<0.05, **p<0.01, ***p<0.001.

### Basal NSCs derive from apical NSCs and are the main contributors to neurogenesis

Several studies have previously shown that the GFAP-expressing NSCs in the V-SVZ give rise to OB interneurons. For example, we have previously shown that most OB adult born interneurons are derived from NSCs in the V-SVZ, which we tagged with a minimal GFAP promoter (Weber *et al*., 2011). However, since the promoter is active in both apical and basal cells, it is not clear whether both pools of NSCs contribute to neurogenesis. To address these points, we selectively tagged apical V-SVZ cells by intraventricular injection of recombinant adeno-associated viral (AAV) particles driving the constitutive expression of GFP in adult WT mice (Luque-Molina *et al*., 2019). We first used this approach to investigate the lineage relationship between apical and basal NSCs. Analysis of coronal brain slices 14 days and 6 weeks after injection showed that at both timepoints the overwhelming majority of GFP^+^ cells were represented by apical cells (supplementary Fig. S6A, B). The rare basal GFP^+^ cells 14 days after the injection were virtually all non-cycling (supplementary Fig. S6B), showing that they were not rapidly proliferating progenitors. Even 6 weeks after injection, GFP^+^ cells displaying NSC markers like Sox9 and/or GFAP were consistently observed at the apical but not the basal side of the V-SVZ (Fig. 5A), confirming that our approach targets apical NSCs. Most apical GFP^+^ cells also expressed Sox9, consistent with previous observations showing the expression of this transcription factor in most ependymal cells (Scott *et al*., 2010; Sun *et al*, 2017).

**Figure 5:**
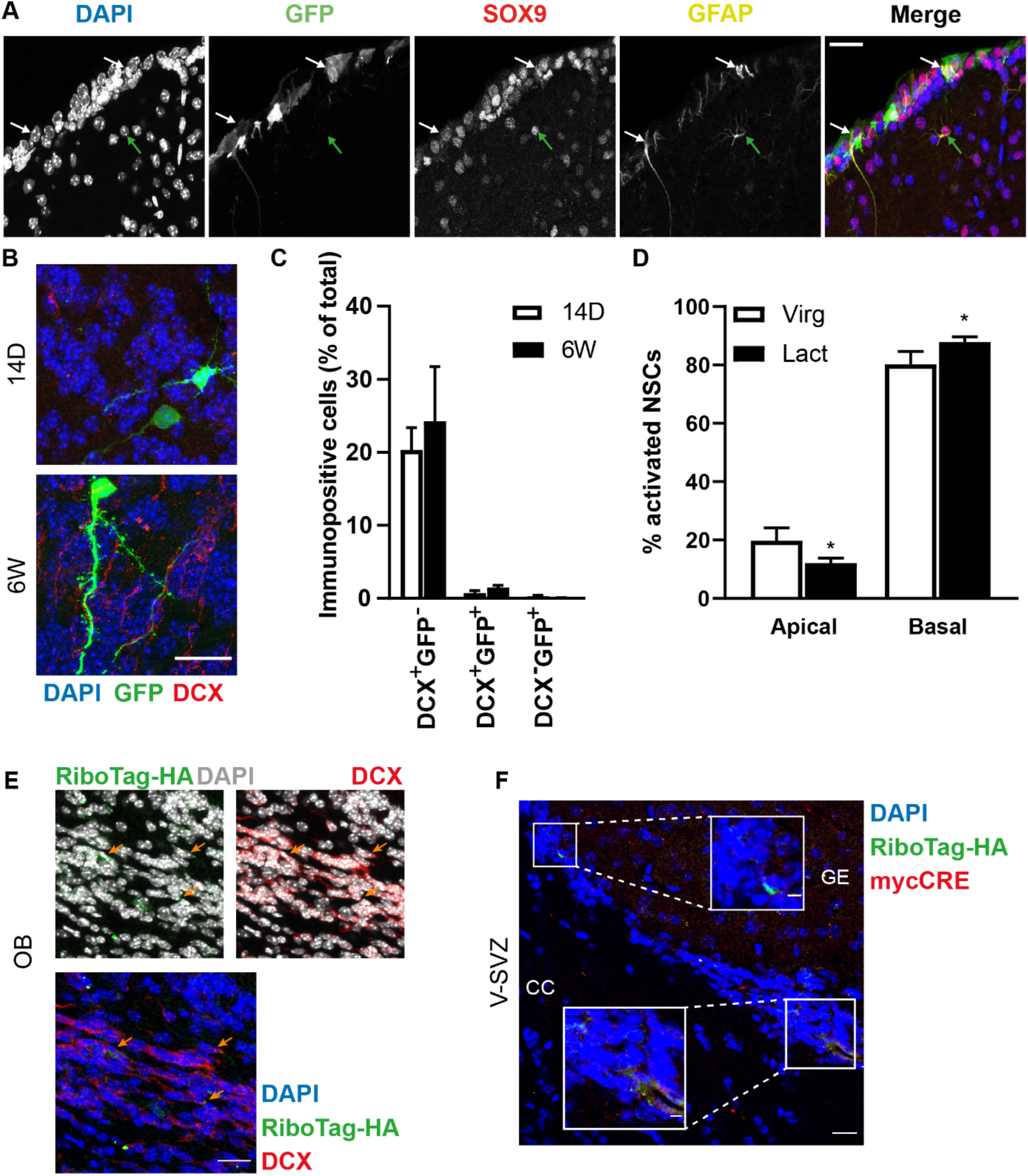
Apical NSCs give rise to few basal NSC-like cells in the SVZ and new OB interneurons. (A) Representative micrographs of coronal sections of WT mice intraventricularly injected with AAV-GFP, and stained for GFAP and Sox9 at 14 days after injection. White and green arrows indicate examples of apical and basal triple positive cells. Scale bar indicates 30 μm. (B) Representative micrographs of the OB of AAV-GFP-injected animals, at 14 days (14D) and 6 weeks (6W) after virus injection. Scale bar indicates 20 μm. (C) Quantification of GFP^+^ and DCX^+^ cells in the OB, as percentage of total DAPI cells. (D) Proportion of apical and basal E^h^ cells counted by flow cytometry and given as percentage of total G^+^ cells from hGFAP;H2B-GFP mice that had been lactating for 7 days (Lact) or virgin littermate controls. Bars represent mean ± SEM.* indicates significance: *p<0.05 N≥5. (E, F) Representative confocal images of coronal slices of the core of the olfactory bulb (OB) (E) and V-SVZ (F) obtained from RiboTag mice 8 weeks after injection of AAV-hGFAP;mycCRE particles in the lateral corner of the V-SVZ. The arrows in (E) indicate Doublecortin (DCX) immunopositive neuroblasts displaying hemagglutinin-tagged ribosomes (RiboTag-HA). In (F) the inset on the left side illustrates cells in the basal V-SVZ co-expressing both myc-CRE and RiboTag-HA, showing activity of the hGFAP promoter as well as recombination. The inset on the right side illustrates a cell at the beginning of the rostromigratory stream expressing only the RiboTag, RiboTag-HA, indicating that the hGFAP promoter was no longer active after recombination.

To investigate whether GFP^+^ cells are involved in neurogenesis, we next quantified doublecortin (DCX) expression and GFP labelling in the OB of injected mice (Fig. 5B, C). At both 14 days and 6 weeks after AAV-GFP injection, we could find three types of labelled cells in the OB: a minority of cells with neuronal morphology expressed GFP only or both DCX and GFP, whereas the vast majority of immature neurons were labelled with only DCX (Fig. 5B, C). Independent of the time of analysis, both types of GFP^+^ cells represented a very small percentage of total cells especially in the core of the OB where newly generated neurons switch migration mode (supplementary Fig. S6C). Taken together, our data show that within a six-week period, apical NSCs generate few basal cells in the V-SVZ with antigenic characteristics of NSCs and rare OB interneurons. However, in normal conditions, the contribution of apical NSCs to the generation of basal NSCs may be difficult to detect due the fact that slow-cycling apical NSCs rarely divide. Therefore, we next investigated the contribution of apical and basal NSCs to the physiological increase in proliferation and consequent OB neurogenesis that is elicited in female mice after seven days of lactation (Shingo *et al*., 2003). For this, we compared the proportion of apical and basal NSCs contributing to the pool of activated NSCs in the V-SVZ in control virgin female hGFAP;H2B-GFP mice and littermate dams, which had been lactating for 7 days (Fig. 5D). This analysis revealed that lactation led to a significant increase in the proportion of activated basal and a parallel decrease the in proportion of activated apical stem cells (Fig. 5D), indicating that basal NSCs mainly contribute to the increased generation of neuronal progenitors. Finally, to directly investigate whether basal NSCs contribute to neurogenesis we injected AAV-hGFAP-mycCRE viral particles to drive the expression of mycCRE under the control of a minimal GFAP promoter in the V-SVZ of RiboTag mice. To selectively tag basal NSCs, we injected mice with either 1×10^6^ or 0.5×10^6^ particles in the basal corner of the V-SVZ and sacrificed the two groups of animals after 5 days and 8 weeks, respectively. Irrespective of time and number of injected viral particles, we found similarly low numbers of tagged cells in the V-SVZ (Fig. 5F and supplementary Fig. S6D). Moreover, the majority of these tagged cells were localized at the dorsal/medial side of the lateral V-SVZ, with a position of the nuclei compatible with their being basal NSCs. Consistent with this, most of these cells not only displayed HA-tagging of the ribosomes, showing that recombination had occurred, but also mycCRE staining (Fig. 5F and supplementary Fig. S6D). However, in mice sacrificed after 8 weeks, cells showing only recombination but not mycCRE staining were also observed at the beginning of the rostral migratory stream (Fig. 5F). Taken together, these observations show that our modified injection strategy allowed us to tag a very small number of cells resident at the basal V-SVZ. Despite the very low number of cells tagged in the V-SVZ, we consistently observed RiboTag-HA-labelled neuroblasts in the core of the OB after 8 weeks. Moreover, tagged neuroblasts represented a subset of the total neuroblasts in the core area of the OB, as would be expected if they derived from the few basal NSCs tagged in the V-SVZ. Taken together, these data show that basal NSCs sustain OB neurogenesis.

### Regulators of NSC activation and notch signalling are differentially expressed in apical and basal NSCs

Upon our reanalysis of the data in Beckervordersandforth *et al*. (2010), we could identify upregulation of the PEDF signaling pathway in activated in Prominin^-^/GFP^+^ cells (supplementary Tables 1, 2) and differential regulation of genes associated with the Notch signaling pathway between the latter and Prominin^+^/GFP^+^ cells (Fig. 6A and supplementary Fig. S7A). Previous studies have revealed that PEDF acts as a niche-derived growth factor promoting NSC self-renewal by upregulating the expression of Notch-signalling effector genes Hes1 and Hes5(Ramirez-Castillejo *et al*., 2006). We have recently found that the TLX/NR2E1-Notch axis coordinates activation and proliferation between apical and basal progenitors in the V-SVZ regulating Notch signalling across the V-SVZ (Luque-Molina *et al*., 2019). However, the molecular basis underlying this regulation is still poorly understood and it is not clear how NSCs contribute to this regulation. To investigate this issue, we used quantitative RT-PCR to compare the expression of *Tlx* and various Notch signaling-related gene transcripts in apical G^+^P^+^E^l^ and basal G^+^P^-^E^l^ quiescent NSCs, in apical G^+^P^+^E^h^ and basal G^+^P^-^E^h^ activated NSCs and in G^-^P^-^E^h^ transit-amplifying progenitors (TAPs; Fig. 6). Consistent with previous studies (Li *et al*, 2012; Obernier *et al*, 2011), we found that the expression of *Tlx* was significantly greater in apical activated G^+^P^+^E^h^ NSCs as well as in G^-^P^-^E^h^ TAPs compared to other cell types. In contrast, quiescent G^+^P^-^E^l^ and activated G^+^P^-^E^h^ basal NSCs did not show a significant regulation of *Tlx* expression (Fig. 6C), indicating that the elevated expression of the orphan nuclear receptor is associated prominently with the activation of apical but not basal NSCs. Analysis of the transcript levels of the Notch ligands, i.e. *Delta1* (*Dll1*; Fig. 6D) and *Jagged 1* (*Jag1*; Fig. 6E), and of the most prominent Notch effector genes in the V-SVZ, i.e. *Hes1* (Fig. 6F) and *Hes5* (Fig. 6G), also revealed different expression profiles between the various apical and basal cell populations. For all genes analysed, including both Notch ligands, the lowest levels of gene expression were observed in apical quiescent G^+^P^+^E^l^ NSCs. Basal quiescent G^+^P^-^E^l^ cells also displayed low levels of *Dll1* expression, but higher expression of *Jag1* than the apical group. Consistent with previous findings (Kawaguchi *et al*, 2013), in both apical and basal NSCs, the expression levels of Notch ligands were positively correlated with NSC activation (Fig. 6D, E). However, whereas in apical NSCs activation was accompanied by an increase in the expression of both Notch ligands, in activated basal NSCs only *Jag1* transcript levels were increased. Additional differences between apical and basal progenitors were observed with respect to the expression of effector genes. The levels of *Hes1* transcripts were upregulated in a comparable manner across the four remaining cell populations, whereas *Hes5* expression was increased only in basal cells, including TAPs (Fig. 6E, F). Taken together, these data show a distinct distribution of genes associated with Notch signalling in NSCs and precursors along the apical/basal axis of the V-SVZ. They also show that at both sides of the niche, NSC activation leads to the acquisition of a transcriptional profile which is characteristic of Notch signal-sending NSCs, whereas quiescent NSCs, especially at the apical side, essentially display a signature of Notch signal-receiving NSCs. Supporting this last conclusion, we also found that Notch signalling activation was stronger in ciliated NSCs and ependymal, which are most apical (supplementary Fig. S7B, C).

**Figure 6:**
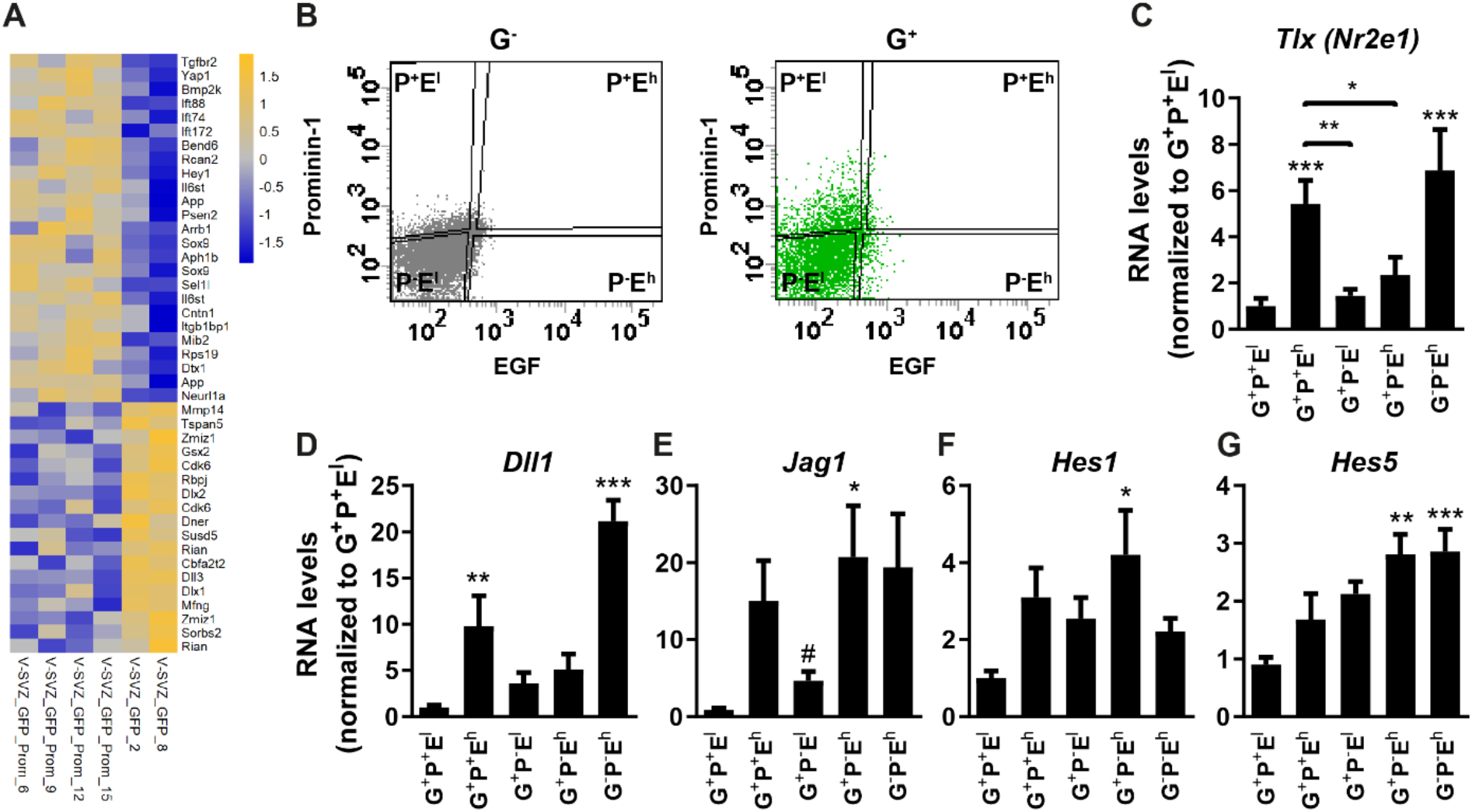
Expression of Notch signaling-related genes in NSCs of the V-SVZ. (A) Heatmap illustrating differentially regulated Notch genes hGFAP;GFP-tagged (GFP) cells expressing Prominin-1 (Prom) as indicated. (B) Representative FACS plots of G^-^ and G^+^ cells sorted for fluorescently labelled EGF and Prominin-1. (C, D) Relative mRNA expression levels for the indicated genes. Bars represent mean ± SEM, N≥4.*/# indicate significance to G^+^P^+^E^l^ population or as indicated (lines), calculated by oneway ANOVA (*) or by Student’s t-test (#). */#p<0.05, **p<0.01, ***p<0.001.

Since the Tlx-Hes1 axis plays a role in the activation of apical NSCs, we next aimed to downregulate *Hes1* expression specifically in apical NSCs via intraventricularly injected AAV constructs, as described before (Luque-Molina *et al*., 2019), to determine the effect on Notch activation and proliferation dynamics in apical and basal NSCs. Using this approach, in addition to apical NSCs, we also target ependymal cells, which also express Hes-1 (Stratton *et al*, 2019). Thus, we first used meta-analysis of the RNA-sequencing data collected and made available by the lab of Sten Linnarsson at Karolinska Institute, Sweden (Zeisel *et al*, 2018), on mousebrain.org to investigate Hes-1 expression in ependymal cells (Supplementary Fig. S8) and to validate our approach. We found that within the ependymal population, the expression of Hes1 was limited to a group of cells half of which were characterized by the expression of Nestin and/or GFAP (Supplementary Fig. S8A, B). Moreover, among ependymal cells, *Hes1*-positive cells population expressed the highest levels of radial glia markers, such as Solute Carrier Family 1 Member 3 (Slc1a3) and Fatty Acid Binding Protein 7 (Fabp7; Supplementary Fig. 87C). Taken together, these data indicate that at the apical side of the niche, Hes1 is expressed mostly in NSCs and less in mature ependymal cells. We therefore proceeded with intraventricular injections of AAVs that drive the constitutive expression of GFP and either a Hes1-specific short hairpin (AAV-Hes1-sh) or a scramble control (AAV-sc-sh) (Luque-Molina *et al*., 2019) to investigate the effect of apical loss of Hes1 function (Fig. 7).

**Figure 7:**
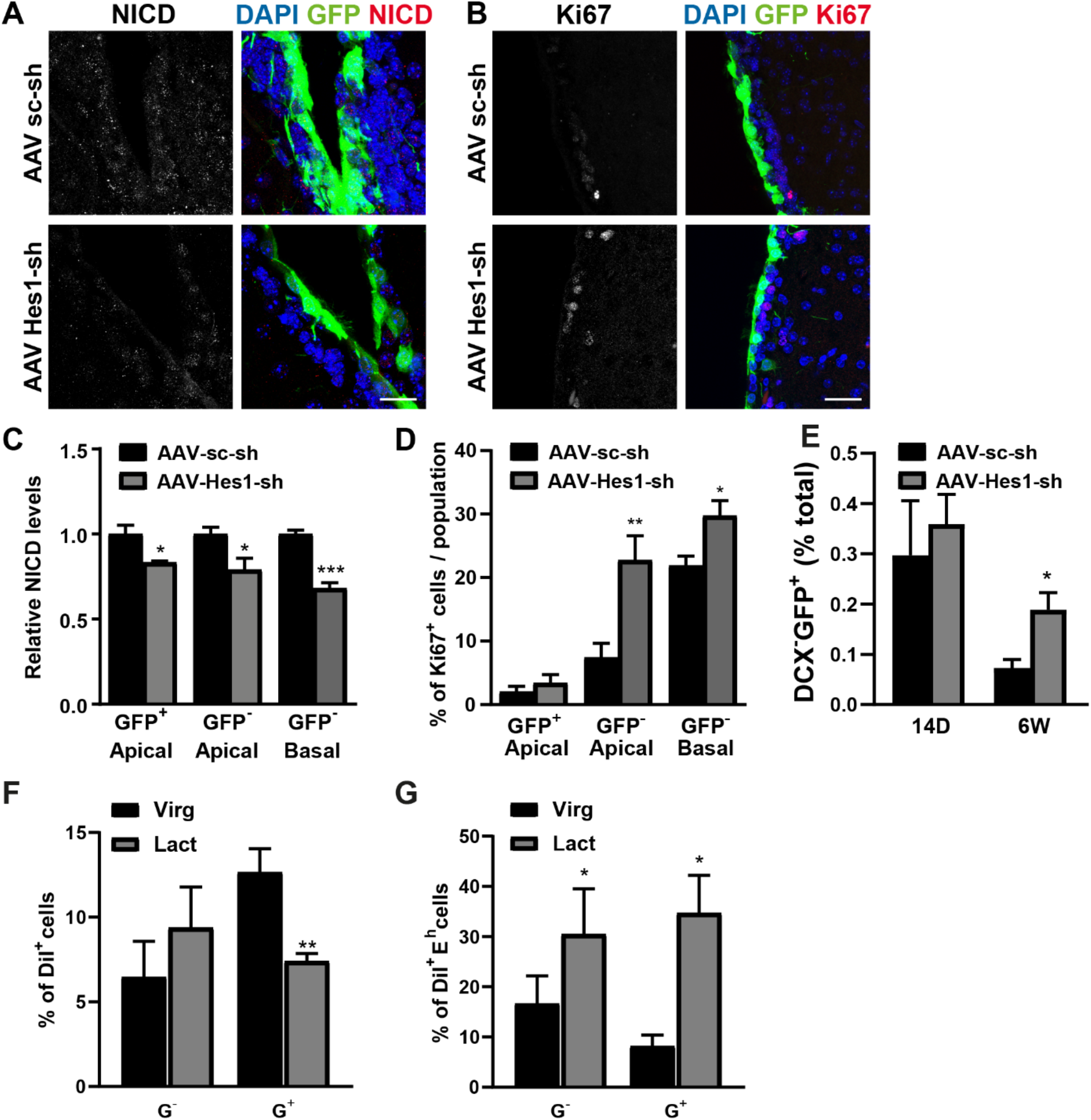
*In vivo* interference with the Notch signalling pathway leads to increased proliferation of apical NSCs. (A, B) Representative confocal photomicrographs the V-SVZs of mice injected with an AAV-expressing GFP and a shRNA against *Hes1* (AAV Hes1-sh) or a scrambled sequence (AAV sc-sh) as a control. Coronal sections were stained with antibodies to either the Notch intracellular domain to visualize cells with activated Notch signaling (NICD, A) or Ki67 (B) to visualize actively cycling cells. DAPI was used for nuclear counterstain. Scale bars = 30 μm. (C, D) Quantification of relative numbers of NICD^+^ (C) or Ki67^+^ cells (D) among infected (GFP^+^) or uninfected (GFP^-^) cells 6 weeks after infection. (E) Quantification of GFP^+^DCX^-^ cells in the olfactory bulb (OB) 14 days (14D) and 6 weeks (6W) after virus injection. (E) GFP+DCX-cells in the olfactory bulb (OB) 14 days (14D) and 6 weeks (6W) after virus injection. (F, G) Quantitative analysis illustrating the total percentage of apical G^+^ and G^-^ cells (F) and the percentage of activated cells in each population after 17 days of lactation (G). Bars represent mean ± SEM, N≥3. * indicates significance within each population. *p<0.05, **p<0.01, ***p<0.001.

Independent of the constructs employed, 14 days after intraventricular AAV injection, GFP was expressed mainly by cells lining the lateral ventricle (Fig. 7A, B). We next measured fluorescence levels of the Notch intracellular domain (NICD; Fig. 7A, C), to quantify activation of Notch signalling, and Ki67 expression (Fig. 7B-D), in order to measure the number of cycling cells in infected (GFP^+^) and non-infected (GFP^-^) at the apical and basal side of the V-SVZ. This analysis showed that, compared to the scramble control, injection of AAV-Hes1-sh affected NICD immunoreactivity across all groups examined including basal cells, showing that Notch signalling connects progenitors between the apical and basal sides of the niche. Downregulation of Notch signalling was accompanied by an increase in the number of cycling cells (Ki67^+^) across all the populations examined, although this effect was stronger in apical than in basal cells (Fig. 7D). Moreover, the increase of proliferation, even at the apical side, was also observed in GFP^-^ non-transduced cells, further underscoring the importance of cell-cell interactions for the regulation of proliferation in the V-SVZ.

Analysis of DCX expression 14 days after AAV injection revealed no effect of of *Hes1* knockdown on the number of neuroblasts (supplementary Fig. S9A). However, AAV-Hes1-sh infection resulted in a significant increase in the number of GFP^+^/DCX^+^ neuroblasts in the apical V-SVZ (supplementary Fig. S9A). At this time, downregulation of *Hes1* expression also led to an increase in the number of GFP^+^ neurons in the OB (Fig. 7E). Strikingly, at both 14 days and 6 weeks after and upon injection of either viral construct, the vast majority of DCX^+^ neuroblasts were represented by GFP^-^ cells. Independent of GFP expression, downregulation of *Hes1* did not affect Sox9 and/or GFAP expression at the apical V-SVZ (supplementary Fig. S9B). Expression of these markers was also not affected in GFP^-^ basal cells (supplementary Fig. S9C), showing that downregulation of *Hes1* does not affect astrocyte generation or NSC self-renewal. However, the number of basal GFP^+^/Sox9^+^ cells was significantly increased after *Hes1* knockdown, suggesting that decreasing Notch signalling may promote loss of apical attachment in NSCs. Taken together, these data show that in the V-SVZ, Notch signalling connects apical and basal progenitors, thereby repressing differentiating cell division across the V-SVZ and promoting neurogenesis and basal relocation in apical NSCs. However, this type of manipulation did not allow us to observe the immediate consequences of downregulation of Notch signalling in apical and basal NSCs. Therefore, we next exposed the whole V-SVZ dissected from 8W-old hGFAP;H2B-GFP mice to the γ-secretase inhibitor DAPT (N-[N-(3,5-difluorophenacetyl)-L-alanyl]-S-phenylglycine t-butyl ester) or the solvent dimethyl sulfoxide (DMSO) as a control, for 24 hours before fixation and immunostaining with antibodies against Nestin and Ki67 (supplementary Fig. S10A, B). The treatment with DAPT resulted in an overall increase in the proportion of Ki67^+^ cells displaying high intensity Ki67 staining (supplementary Fig. S10A, C) and nucler morphology of cells in meta- or anaphase, (supplementary Fig. D). Blockade of Notch signaling also led to a significant increase in the percentage of G^+^Nestin^+^ and G^+^Ki67^+^Nestin^+^ triple positive cells in the G^+^ population (supplementary Fig. S10E, F). Notably, these effects were only significant at the apical and not at the basal side of the niche, supporting the view that inhibition of Notch signaling primarily affects apical NSCs.

It has been shown that increasing progenitor numbers in the V-SVZ decreases the proliferation of NSCs. Moreover, it was shown that this effect involves downregulation of Notch-lateral inhibition and Notch activation in NSCs(Aguirre *et al*, 2010). We therefore next investigated whether this regulation affects both apical and basal NSCs. For this we determined the number of total apical and basal NSCs and the percentage of activated NSCs 14 and 17 days after lactation. The only significant change detected, was a decreased in the number of apical NSCs after 17 days of lactation (Fig. 7F). Moreover, at this time the number of activated apical NSCs was also increased, consistent with a decrease in Notch signalling in the apical V-SVZ (Fig. 7G). Taken together, these data indicate a scenario in which apical but not basal NSCs respond to increased progenitor proliferation in the V-SVZ by decreasing Notch signalling and proliferation.

## Discussion

In this study we show the presence of basal NSCs in the adult V-SVZ of the mouse brain for the first time. Notably, our data show that basal NSCs are responsible for the generation of most adult-born interneurons in the OB. Apical NSCs, in contrast, regulate the pace of differentiating cell division across the V-SVZ by instructing the strength of Notch activation across the niche, thereby acting as gate-keepers for neurogenesis. It is well established that Notch signalling is essential to maintain stemness during quiescence (Imayoshi *et al*, 2010; Kawaguchi *et al*., 2013; Kawai *et al*, 2017; Than-Trong *et al*, 2018). It is also known that interaction between Notch and EGFR signalling control niche homeostasis by coordinating the proliferation of adult NSCs and more differentiated progenitors (Aguirre *et al*., 2010; Wang *et al*, 2009). However, the mechanisms underlying this regulation are still not clear as they were investigated in a situation of permanent upregulation of EGFR signalling. We show that these function of Notch signalling are based on a directional distribution of Notch ligands which increase not only according to lineage progression, as reported before (Kawaguchi *et al*., 2013), but also along the apical/basal axis of the V-SVZ, thereby providing apical NSCs with essential characteristics of Notch receiving cells. Moreover, our data indicate that different modulators of Notch signalling play a role in apical and basal NSCs. Whereas PEDF seems to regulate Notch signalling in basal NSCs, apical NSCs are more responsive to the change in the number of progenitors (Aguirre *et al*., 2010). We have further analysed the relationship between NSC proliferation and progenitor number in the context of a transitory increase in the number of proliferating progenitors following lactation. In light of the characteristics of Notch receiving cells, we propose that the increase in progenitor numbers leads to an increase in Notch activation prevalently in apical NSCs and to a downregulation of their proliferation. Upon differentiation and migration of the extra-proliferating progenitors outside of the V-SVZ, Notch signalling in apical NSCs will normalize leading to a consequent upregulation of EGFR signalling in apical NSCs.

The identity of NSCs in the V-SVZ has been object of numerous studies. Despite past disagreements on the nature of NSCs, according to the general consensus, apical cells are considered the main source of OB neurogenesis (Beckervordersandforth *et al*., 2010; Codega *et al*., 2014; Doetsch *et al*., 1999; Johansson *et al*, 1999; Nam & Capecchi, 2020). In contrast with this view, we here discovered the presence of basal, non-radial-glia-like NSCs in the adult V-SVZ. Previous work of transcriptional characterization of cells in the adult V-SVZ has also provided evidence for the presence of different pools of NSCs (Dulken *et al*, 2017; Luo *et al*, 2015; Mich *et al*, 2014; Morizur *et al*, 2018). However, this work did not investigate the localization of the various NSC pools in the niche, associating the various NSC types essentially with different degrees of quiescence and activation. Thus, our finding that basal and apical NSCs display differences in cell cycle progression provides additional insight for the interpretation of these previous data. Moreover, it also provides a more unified view of the spatial organization of different pools of adult NSCs in the two main neurogenic niches. The presence of morphologically distinct types of NSCs, i.e. type 1 and type 2 (Steiner *et al*, 2006), has been long studied in the in the SGZ. Although both pools of hippocampal NSCs express radial glial markers in addition to Sox2 and Sox9 transcription factors (Suh *et al*, 2007; Sun *et al*., 2017), they are morphologically distinct, as only type 1 cells display a typical radial glia morphology with apical-basal polarity (Seri *et al*, 2001). In striking similarity with the differences between apical and basal NSCs in the V-SVZ reported here, it was observed that in the SGZ, type 1 radial NSCs consistently display Nestin and GFAP immunoreactivity although they are largely quiescent (Sun *et al*., 2017). In fact, in the adult SGZ, the expression of proliferation markers was mainly observed in non-radial NSCs, which like basal NSCs in the V-SVZ are the main contributors to steady-state neurogenesis and can actively proliferate in response to physiological stimuli (Lugert *et al*, 2010).

Although generally considered a NSC marker, Nestin immunoreactivity is not always observed in NSCs, especially if they are undergoing quiescence(Codega *et al*., 2014). Indeed, we here found here that Nestin expression is significantly downregulated with increasing age and in quiescent apical NSCs. However, in adult basal NSCs, Nestin expression is virtually absent, independent of the state of activation, suggesting that the expression of the intermediate filament is not only a function of the cell cycle state. Indeed, in the SGZ, Nestin is also expressed in radial NSCs, which are largely quiescent (Encinas *et al*, 2006; Kempermann *et al*, 2004). Notably, unlike in the V-SVZ, NSCs and progenitors in the SGZ all display primary cilia (Amador-Arjona *et al*, 2011; Breunig *et al*, 2008). Since we here found that the majority of ciliated apical NSCs extend primary cilia, it is possible that cilia-dependent signals promote Nestin expression. In the V-SVZ, primary cilia do not only regulate quiescence in radial glia and adult NSCs (Beckervordersandforth *et al*., 2010; Khatri *et al*., 2014), but they are also key to sensing signals present in the cerebrospinal fluid filling the ventricular cavity (Monaco *et al*., 2019; Silva-Vargas *et al*, 2016). Consistent with this view, recent observations have highlighted the possibility that in the V-SVZ, acquisition of quiescence occurs over a prolonged time period (Borrett *et al*, 2020) and involves different stages as well as the integration of multiple regulatory signals.

Thus, the postnatal V-SVZ contains two pools of NSCs that are exposed to different microenvironments and are differentially equipped to sense and transduce niche signals.

## Material and Methods

### Animals and V-SVZ dissection

All animal experiments were approved by the Regierungspräsidium Karlsruhe and the local authorities of Heidelberg University. For FACS, C57BL/6 (Wildtype, WT) and hGFAP-tTA;H2B-GFP (hGFAP;H2B-GFP) mice were sacrificed by CO_2_ inhalation followed by cervical dislocation (adult), or by decapitation (neonatal). For DiI (1,1’-Dioctadecyl-3,3,3’,3’-Tetramethylindocarbocyanine Perchlorate, ThermoFisher Scientific) analyses, brains were placed on a petri dish. After incision of the hemispheres the exposed apical surface of the V-SVZ was labelled by applying DiI, dissolved in ethanol, for 1 minute followed by washing in sort medium. Thereafter the V-SVZ was dissected and further analysed as described below. To analyse the effect of lactation, hGFAP;H2B-GFP pregnant dams were sacrificed between seven and seventeen days after having given birth. Littermate virgin controls were sacrificed at the same time. For immunofluorescence WT, hGFAP;H2B-GFP and RiboTag mice (Sanz *et al*, 2009) were sacrificed by intracardial perfusion as described below.

### Fluorescence activated cell sorting (FACS)

V-SVZ cells were dissociated and incubated at 4°C in sort medium containing anti-Prominin1-APC, anti-Prominin-1-PE or anti-Prominin-1-BV421 for 30 min. For experiments requiring EGFR labelling, cells were incubated for a further 30 min with recombinant human EGF conjugated to Alexa Fluor 488 or Alexa Fluor 647. Thereafter, cells were rinsed twice in sort medium before being stained with Propidium Iodide (PI, 1:100) to stain dead cells and sorted with a FACSAria III flow cytometer (BD Biosciences, Heidelberg, Germany) as previously described (Cesetti *et al*., 2009; Ciccolini *et al*., 2005; Khatri *et al*., 2014). For clonal analysis, FACS sorted neural stem cells were plated for neurosphere formation as previously explained (Ciccolini *et al*., 2005).

For the analysis of self-renewal *in vivo*, we calculated the percentage of H2B-GFP expressing (G^+^) cells displaying high or intermediate levels of fluorescence as described before (Luque-Molina *et al*., 2017). Briefly, H2B-GFP fluorescence levels were plotted on a logarithmic scale between the values of 10^2^ and 10^5^. Based on negative controls, fluorescence levels around 10^2^, 10^3^, and 10^4^-10^5^ were considered negative, intermediate, and high, respectively.

For a detailed list of antibodies, please see supplementary table S3.

### RNA expression analysis and qPCR

RNA was isolated with Arcturus^®^ PicoPure™ RNA Isolation Kit (ThermoFisher Scientific) according to manufacturer’s protocol including DNase digestion. Cells were directly sorted into 100 μl Lysis buffer, vortexed and stored at −80°C until RNA isolation. For reverse transcription, 9 μl purified RNA was incubated with 1 μl Oligo(dT)15 primers (Promega) at 80°C for 3 min. M-MLV buffer, M-MLV reverse transcriptase enzyme (200 U/ μl, Promega), dNTPs (10mM) and RNase-free water were added to RNA and Oligo(dT)15 mix to reach a final volume of 20 μl. cDNA synthesis was performed at 42°C for 60 min followed by 10 min at 80°C. cDNA was then preserved at −20°C until qPCR. mRNA levels for the genes of interest were measured using a StepOnePlus Real-time PCR system and TaqMan gene expression assays (Applied Biosystems): β-actin (ID: Mm01205647_g1), Tlx (ID: Mm00455855_m1), Hes1 (ID: Mm01342805_m), Hes5 (ID: Mm00439311_g1), Dll1 (ID: Mm01279269_m1), Jag1 (ID: Mm01270195_gh), Lrig1 (ID: Mm00456116_m1). Cycle threshold (Ct) values were each normalized to the respective β-actin value (= ΔCt). ΔΔCt was calculated by subtracting the ΔCt value of each population from the ΔCt of the G^+^P^+^E^-^ population (normalizing population).

### Intraventricular injections

WT mice were injected in the lateral ventricle with 1 μL of adeno-associated virus (AAV) with a plasmid encoding short hairpin RNA targeting *Hes1* under the control of a U6 promoter (AAV-Hes1-sh), or a scramble control sequence (AAV-sc-sh). Characteristics of the plasmid and procedures of injection were described previously (Luque-Molina *et al*., 2019). RiboTag mice were injected in the lateral corner of the V-SVZ with 1×10^6^ or 0.5×10^6^ of AVV-hGFAP;myc-CRE viral particles to drive the expression of a myc-tagged constitutively active CRE recombinase under the control of a human GFAP promoter (Tang *et al*, 2015). Mice were sacrificed and processed for immunostaining 5 days and 8 weeks after injections as described below.

### Immunofluorescence

Whole mount dissection and immunostaining was performed as previously described (Mirzadeh *et al*., 2008; Monaco *et al*., 2019).

For coronal sections, mice sedated with pentobarbital (400 mg/kg) were perfused with ice-cold phosphate-buffered saline (PBS) followed by 4% paraformaldehyde (PFA) in PBS, after which the brains were removed and postfixed in 3% PFA / 4% sucrose in PBS for 48h at 4°C. Brains were then cryoprotected by submersion in 30% sucrose in PBS at 4°C for 24h and sliced into 20-30 μm coronal sections using a Leica CM1950 cryostat (Leica Microsystems, Wetzlar, Germany). The sections were stored in PBS containing 0.01% sodium azide at 4°C until immunostaining. Slices were permeabilized with 0.5% NP-40 in PBS for 5 min, then incubated in 10 mM glycine for 30 min for reduction of background fluorescence. Slices were blocked with 5% fetal calf serum (FCS) in PBS for 1h, and incubated with primary antibodies in PBS at 4°C overnight. After washing, secondary antibodies were applied in 5% FCS in PBS containing 4’,6-diamidino-2-phenylindole (DAPI) for nuclear counterstain for 2h. Slices were then washed and mounted in Mowiol. All steps were performed at room temperature if not otherwise indicated. Slices and whole mounts that were to be directly compared in terms of protein expression and fluorescence level were stained in parallel using the same antibodies and dilutions. Immunofluorescence on sorted cells was performed as previously described (Khatri *et al*., 2014). A detailed list of primary antibodies used is provided in supplementary table S4.

### Imaging

Coronal sections and whole mounts were imaged using a Leica SP8 laser scanning confocal microscope. Images were acquired as z-stacks with 1 μm steps or 0.5-0.7 μm steps for cilia imaging. The imaging settings (laser intensity, gain, offset) were kept consistent for each experiment to ensure comparability.

### Transcriptome analysis by microarray

Transcriptome analysis was performed on the RNA microarray dataset from (Beckervordersandforth *et al*., 2010), available on the Gene Expression Omnibus (GEO) database (http://www.ncbi.nlm.nih.gov/gds), accession number GSE18765. Briefly, ependymal and GFP expressing cells, coexpressing Prominin-1 or not, were isolated by FACS from the V-SVZ and the diencephalon of hGFAP-eGFP mice. Expression was analysed after hybridization of total RNA from from each cell group on Affymetrix MOE430 2.0 arrays. GFP^+^Prom^+^ and GFP^+^ samples were normalized with the Transcriptome Analysis Console (TAC; version 4.0.1.36; Thermo Fisher Scientific) using standard RMA settings. TAC was also used for the PCA of all samples. Statistical analysis was done as described in Beckervordersandforth *et al*. (2010) and genes with a raw p-value < 0.05 were used to define the set of 4713 regulated genes. Additional filters for fold-change > 1.3x and linear average expression > 100 in at least one group were applied. Heatmaps were generated in R. Pathway analyses were done through the use of QIAGEN’s Ingenuity Pathway Analysis (IPA^®^, QIAGEN Redwood City, www.qiagen.com/ingenuity) using Fisher’s Exact Test p-values. Genes involved in Notch signalling were obtained from Gene Ontology (GO:0007219).

### Statistical analysis

Immunopositive cells were counted and normalized to the total number of cells (identified by DAPI-stained nuclei) or as stated otherwise. For quantification of cells in the V-SVZ, we defined cells as apical if their nuclei were located within the first two rows of nuclei bordering the lateral ventricle. Three regions of each slice were analysed in a 30,000 μm^2^-sized picture aligned with the longest axis of the apical side of the V-SVZ: dorsal, medial and ventral. Fiji/ImageJ software (Schindelin *et al*, 2012) was used for fluorescence intensity measurement and normalization to background levels.

During analysis, the pictures were labelled by treatment (i.e. doxycycline, DAPT, scramble/Hes1 shRNA, pregnancy) at all times and were not randomized or blinded. Statistical significance was determined by two-tailed homoscedastic (or paired, when appropriate) Student’s t-test for comparing two, and one-way or two-way ANOVA with Tukey’s multiple comparisons test for comparing more than two datasets. The analyses were performed using GraphPad Prism software. Significance was reached for * *p*<0.05, ** *p*<0.01, and *** *p*<0.001. Graphs represent mean values ± SEM.

## Acknowledgements

K.B. was supported by the Interdisciplinary Center for Neuroscience (IZN) and the Landesgraduiertenförderung (LGF) of the Heidelberg University Graduate Academy. The authors would like to thank Inma Luque-Molina for her help with clonal analysis. We would also like to thank the Department of Neurobiology for its continued support.

## Author contributions

Conceptualization, Methodology, Funding Acquisition: F.C.; Formal analysis: K.B., Y.A. M.I. and F.C.; Investigation: K.B., Y.A., C.M., G.H.-W., Y.S., U.E., P.K., A.M.H and F.C.; Writing - original draft: F.C., K.B. and Y.A.; Writing - review and editing: F.C., K.B., Y.A., Y.S., U.E., P.K., A.M.H., M.I. and J.B.; Visualization: K.B. and Y.A.

