## supplementary figure and text for "A novel stem cell type at the basal side of the subventricular zone maintains adult neurogenesis"

**Supplementary Table S1:** Selected significantly ($p < 0.05$) enriched canonical pathways in $Prom^{-}GFP^{+}$ cells compared to $GFP^{+}Prom^{+}$ cells. Z-scores > 2 indicate significant activation of the respective pathway.

| ***Pathway*** | **-log(p-value)** | **Ratio** | **z-score** |
|---:|:---:|:---:|:---:|
| *Cell Cycle Control of Chromosomal Replication* | 3.07 | 0.30 | 3.64 |
| *PEDF Signaling* | 1.80 | 0.23 | 3.15 |
| *Sirtuin Signaling Pathway* | 5.39 | 0.23 | 2.99 |
| *Aryl Hydrocarbon Receptor Signaling* | 2.12 | 0.21 | 2.60 |
| *Mitotic Roles of Polo-Like Kinase* | 1.87 | 0.24 | 2.53 |
| *Death Receptor Signaling* | 2.07 | 0.23 | 2.13 |
| *Inhibition of Angiogenesis by TSP1* | 2.40 | 0.32 | 2.12 |
| *EGF Signaling* | 2.70 | 0.29 | 1.81 |
| *Kinetochore Metaphase Signaling Pathway* | 7.27 | 0.34 | 1.67 |
| *Estrogen-mediated S-phase Entry* | 4.24 | 0.46 | 1.51 |
| *FGF Signaling* | 2.46 | 0.25 | 1.34 |
| *Autophagy* | 1.35 | 0.18 | 1.30 |
| *ERK/MAPK Signaling* | 2.31 | 0.20 | 1.12 |
| *Cyclins and Cell Cycle Regulation* | 4.08 | 0.30 | 1.09 |
| *PDGF Signaling* | 2.33 | 0.24 | 0.89 |

**Supplementary Table S3:** Antibodies and recombinant proteins used for flow cytometry.

| Antibody | Conjugate | Host | Company, Catalog # | Lot # | Concentration |
|---|---|---|---|---|---|
| Prominin-1 | APC | rat | Miltenyi Biotec, 130-102-197 | 5190517507 | 1:100 |
| Prominin-1 | PE | rat | Miltenyi Biotec, 130-102-210 | 5181126230 | 1:100 |
| Prominin-1 | BV421 | rat | BioLegend, 141213 | B255870 | 1:1000 |
| EGF | 488 | N/A | Invitrogen, E13345 | 1802775 | 1:1000 |
| EGF | 647 | N/A | Invitrogen, 35351 | 1700382 | 1:1000 |

**Supplementary Table S4:** Antibodies used for immunofluorescence.

| Antibody | Host | Company, Catalog # | Lot # | Concentration |
|---|---|---|---|---|
| Acetylated Tubulin | mouse | Sigma Aldrich, T6793 | N/A | 1:1000 |
| Adcy3 | rabbit | ThermoFisher, PA5-35382 | UL2902981 | 1:500 |
| DCX | mouse | Santa Cruz, sc-271390 | D2720 | 1:200 |
| GFAP | rabbit | Agilent, Z0334-01 | 00059585 | 1:500 |
| GFP | rabbit | Molecular Probes, AA11122 | 1828014 | 1:500 |
| HA | rabbit | Cell Signalling, 3724 | N/A | 1:1000 |
| Ki67 | rabbit | Abcam, 16667 | GR3313195-18 | 1:100 |
| Nestin | mouse | BD Pharmingen, 556309 | 8135889 | 1:100 |
| NICD | rabbit | Abcam, ab8925 | GR218543-39 | 1:2000 |
| Myc | mouse | Santa Cruz, sc-40 | N/A | 1:250 |
| Sox9 | rabbit | Millipore, AB5535 | 3191390 | 1:2000 |

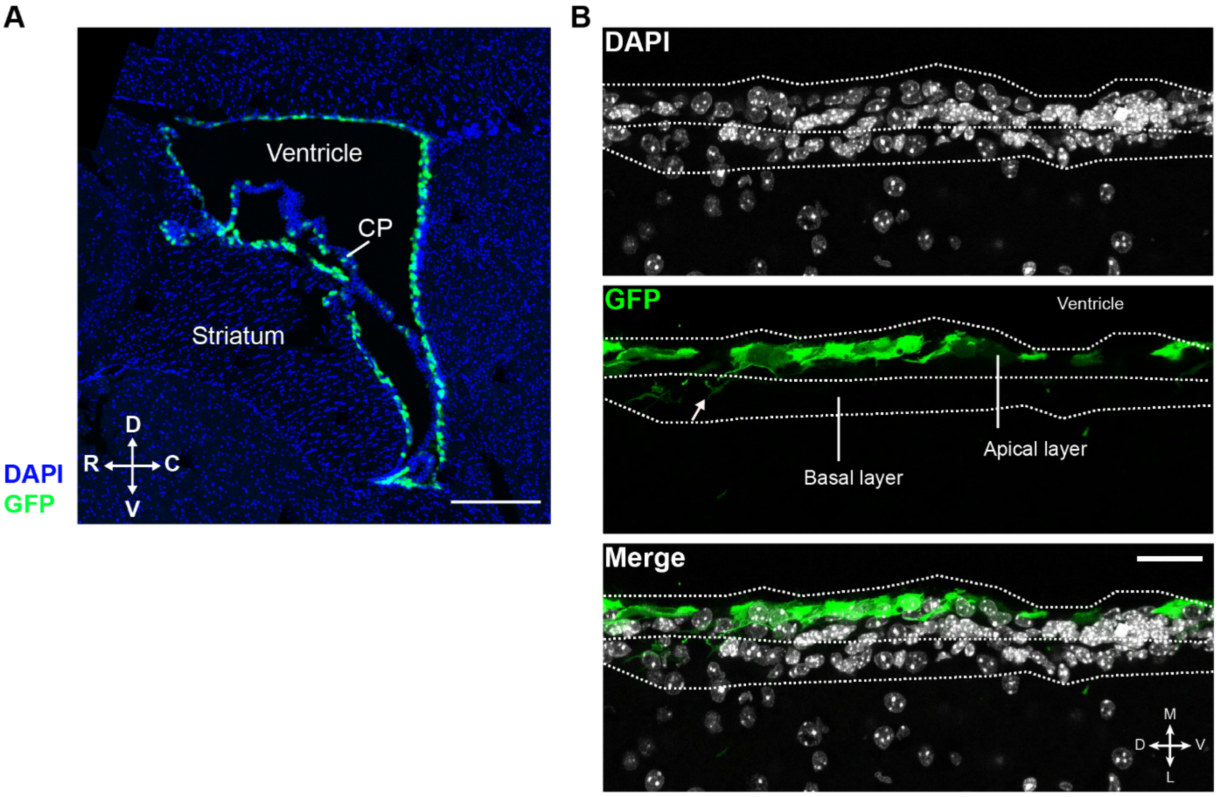


**Figure S1: Apical and basal cells in the V-SVZ.** (A) Representative micrograph of the V-SVZ in a sagittal brain section. The apical surface of the V-SVZ was labelled by injection of AAV-sc-sh viral particles in the ventricular space. Scale bar = 1 mm, CP = choroid plexus, C = caudal, D = dorsal, R = rostral, V = ventral. (B) Detail micrograph of infected cells in a coronal brain section. D = dorsal, L = lateral, M = medial, V = ventral. Scale bar = 30 µm.

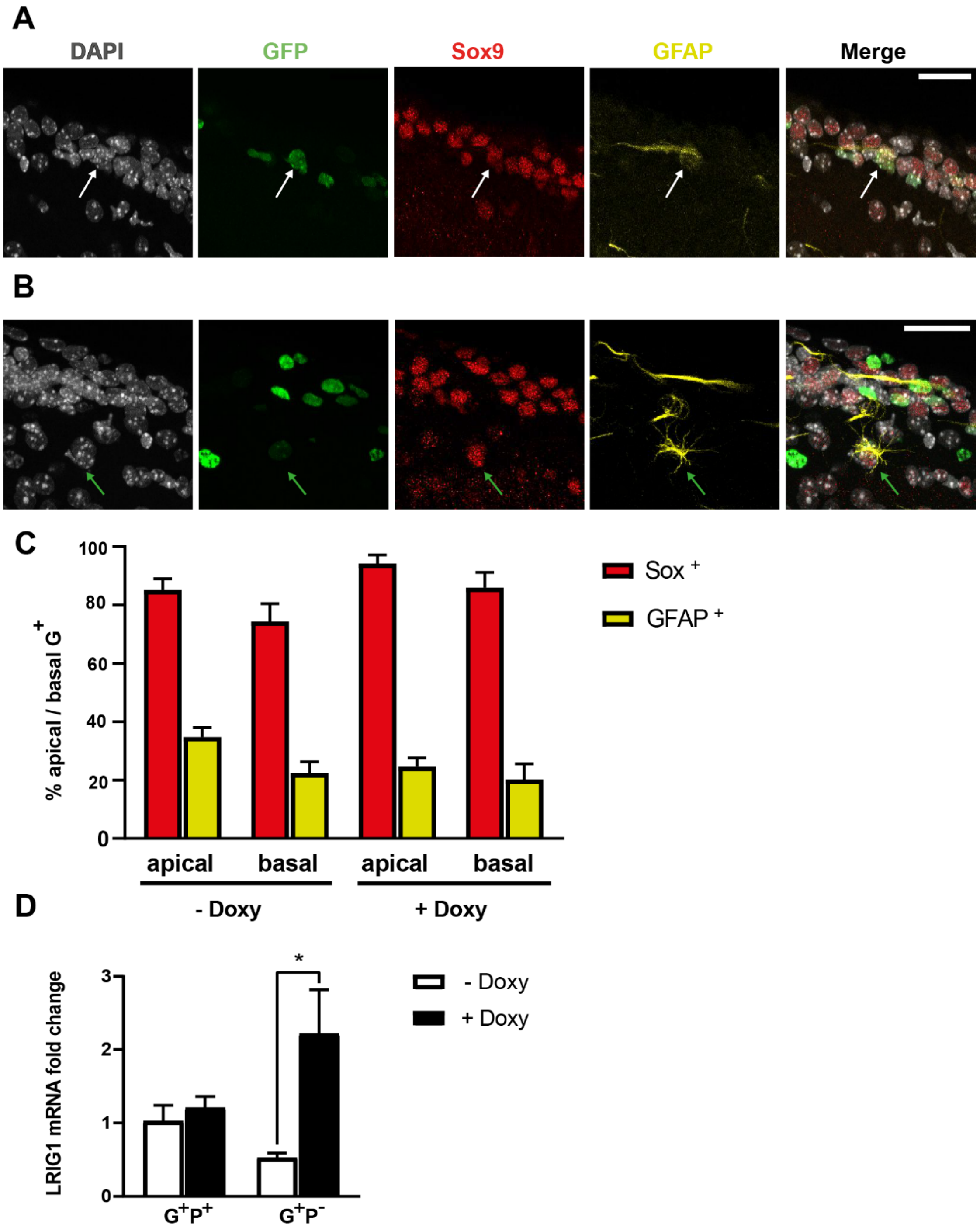


**Figure S2: Basal NSCs in the SVZ express less GFAP than apical NSCs.** (A,B) Representative confocal microphotographs of coronal sections from 8w hGFAP-H2B.GFP mice, immunostained for GFAP and Sox9. White and green arrows point at triple positive apical and basal cells, respectively. Scale bar indicates 20 µm. (C) Quantification of $G^+$ NSCs co-expressing GFAP and Sox9. (D) Expression of *Lrig1* mRNA in untreated (- Doxy) and doxycycline-treated (+ Doxy) apical ($P^+$)and basal ($P^-$) $G^+$ NSCs. Bars represent mean ± SEM. *p<0.05.

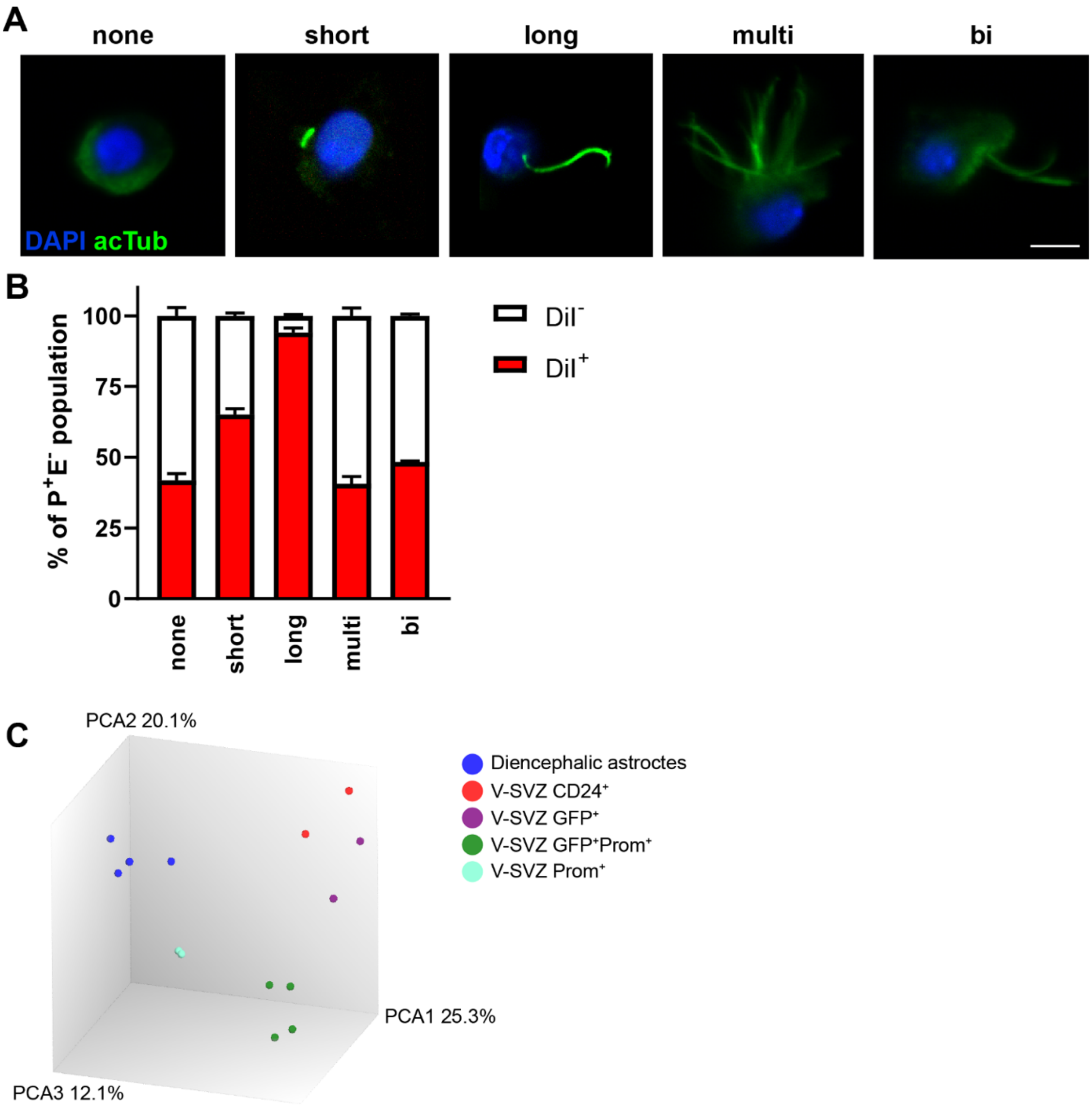


**Figure S3: Different ciliation in apical (DiI$^+$) and basal (DiI$^-$) FACS sorted cells.** (A) Representative microphotographs of FACS-sorted cells immunostained for acetylated tubulin (acTub) and counterstained with nuclear DAPI. Cells are classified as not ciliated (none), short single ciliated (short), long single ciliated (long), multi-ciliated (more than two cilia, multi), or bi-ciliated (two cilia, bi). Sale bar indicates 10 µm. (B) Quantification of P$^+$E$^-$ cells according to ciliation and DiI-staining. Bars represent mean ± SEM. N=5. C) Principal component analysis referring to microarray analysis illustrated in Beckervordersandforth et al. 2010.

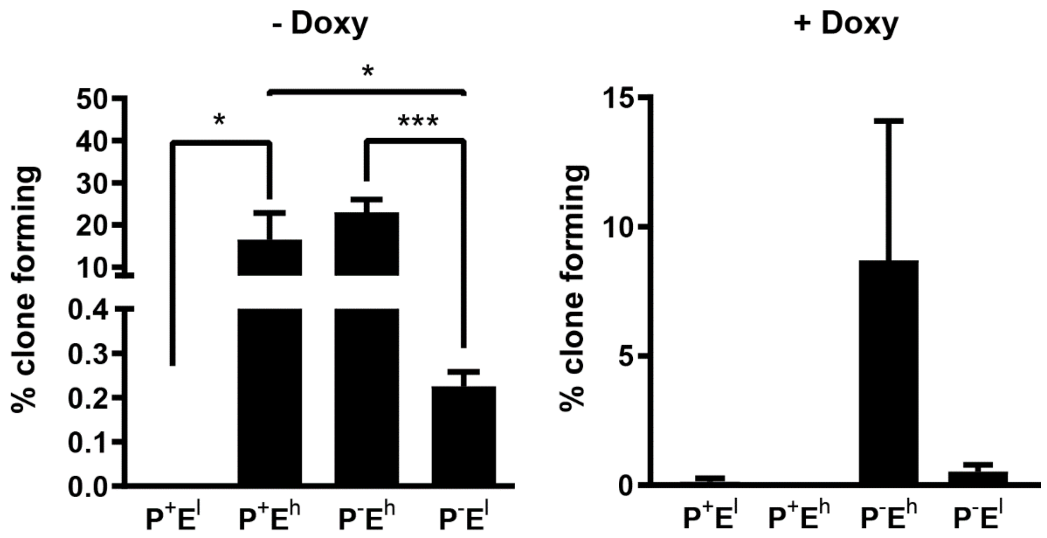


**Figure S4: Label-retaining cells show less clonogenic potential.** $G^+$ cells of untreated (- Doxy) and doxycycline-treated animals (+ Doxy) were labelled for Prominin-1 and EGFR, sorted and number of clone-forming cells analysed. Bars represent mean ± SEM. N=6 (- Doxy), N=5 (+ Doxy)**.** *$p$<0.05, ***$p$<0.001**.**

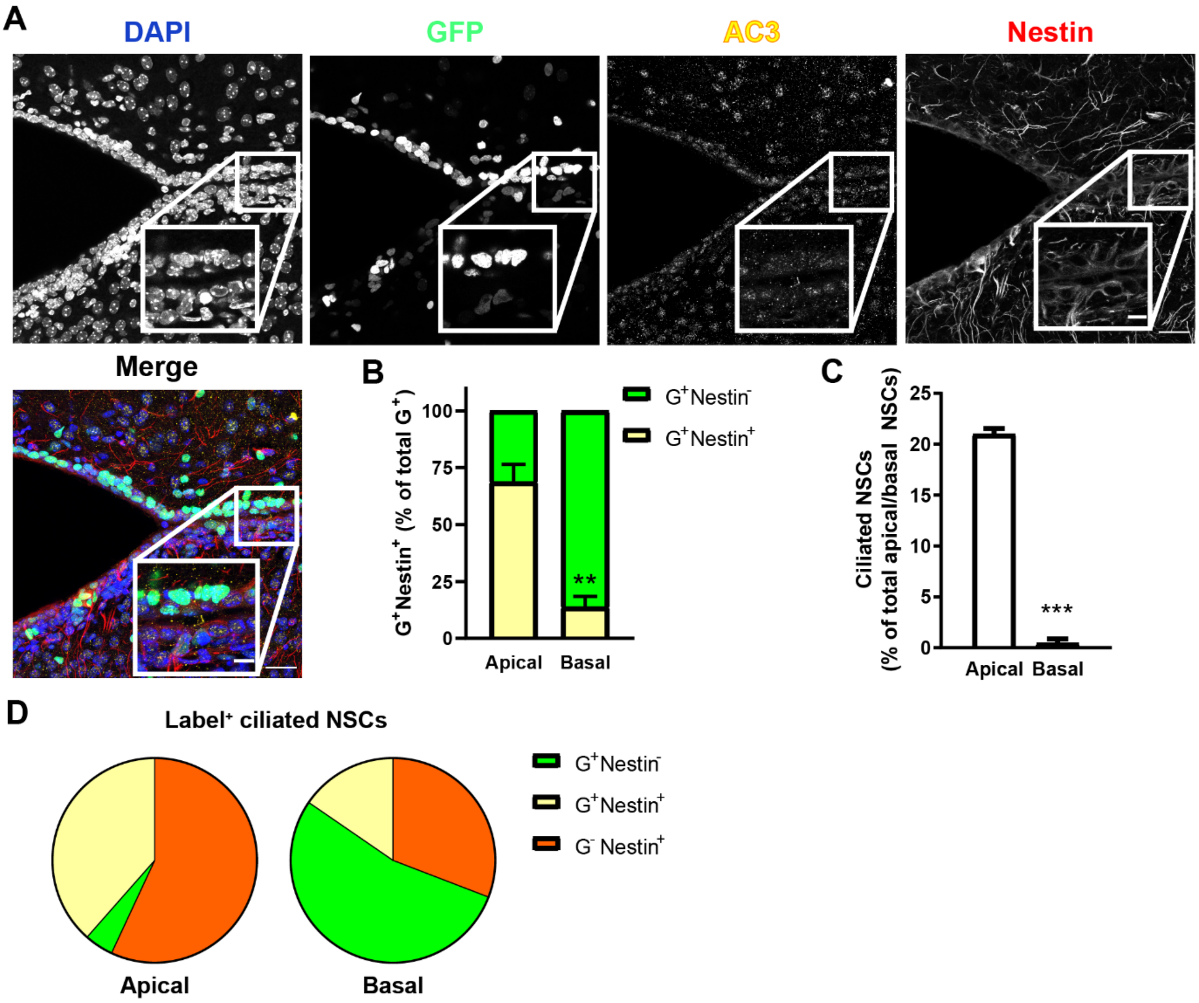


**Figure S5: Expression of H2B-GFP, Nestin and primary cilia in the postnatal SVZ.** (A) Representative confocal microphotographs illustrating immunofluorescent staining of coronal slices from postnatal day 7 (P7) hGFAP-tTA;H2B-GFP mice without doxycycline treatment, with the stem cell marker Nestin (red) and the primary cilia marker adenylate cyclase 3 (AC3, yellow). DAPI was used for nuclear counterstain. Scale bars indicate 30 µm, 10 µm for inserts. (B) Quantification of marker expression in apical and basal cells. (C) Quantification of the proportion of ciliated $G^+$ NSCs in the apical and basal cell layer according to stem cell marker. (D) Pie charts represent antigenic characteristics of apical and basal ciliated cells. Quantification of antigenic characteristics of ciliated apical and basal NSCs. Ciliated, non-labelled cells were disregarded. Bars represent mean ± SEM. * indicates significance between apical and basal cell populations. **$p<0.01$, ***$p<0.001$. N=4.

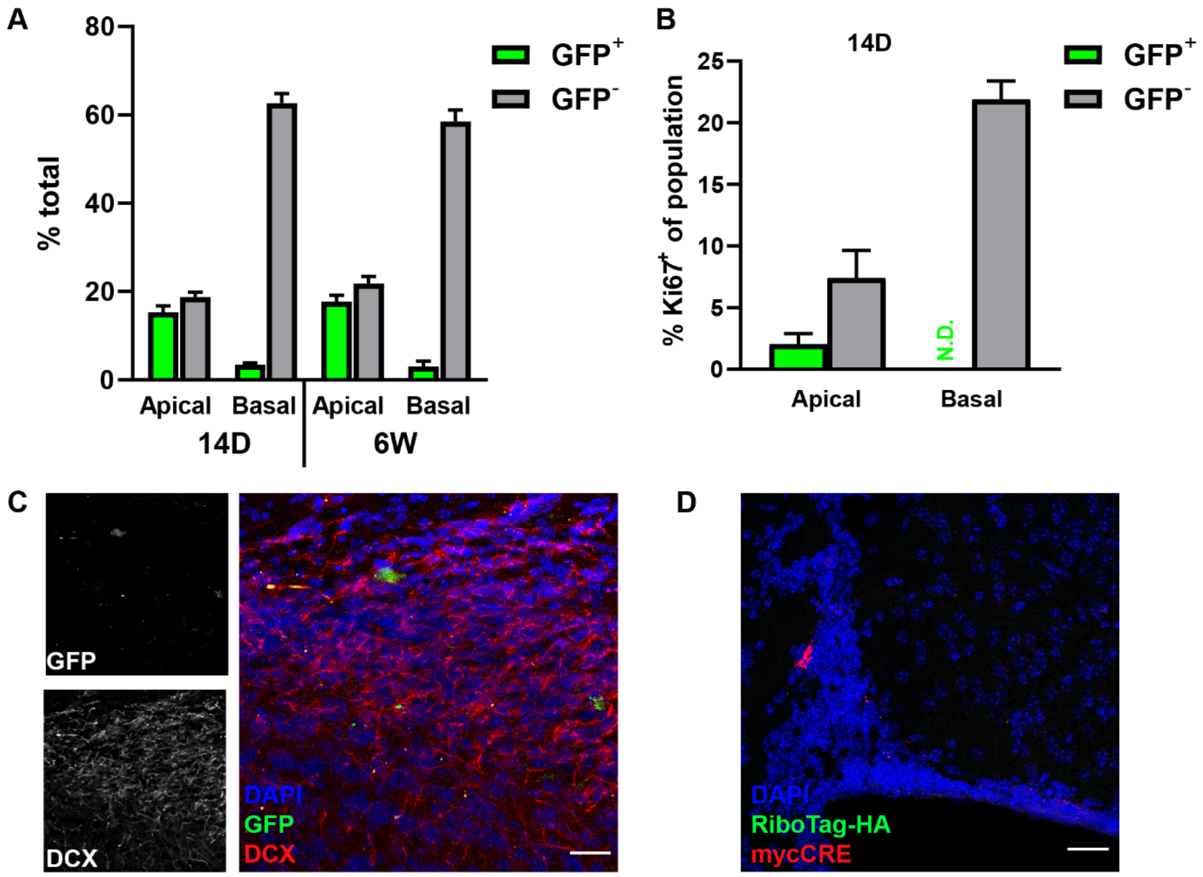


**Figure S6: GFP-infected cells in the V-SVZ.** (A) Quantification of AAV-GFP infected ($GFP^+$) and non-infected cells ($GFP^-$) in the apical and basal layer of the V-SVZ at 14 days (14D) and 6 weeks (6W) post injection into the lateral ventricle, as percentage of total cells. (B) Quantification of AAV-infected or non-infected cells expressing Ki67 in the apical and basal cell layer 14 days after injection into the lateral ventricle. N.D. = none detected. (C) Representative micrograph of newborn neurons in the OB 6 weeks after intraventricular AAV-GFP injection, stained with DCX. Scale bar indicates 20 µm. Bars represent mean ± SEM. N=5. (D) Representative confocal images of coronal slices of obtained from RiboTag mice 5 days after injection of AVV-hGFAP-mycCRE particles in the lateral corner of the V-SVZ. Panels illustrate Doublecortin a few cells expressing mycCRE around the lateral ventricle, showing activity of the hGFAP promoter. Note at this stage the vast majority of the cells do not display hemagglutinin-tagged ribosomes (RiboTag-HA), indicating lack of recombination.

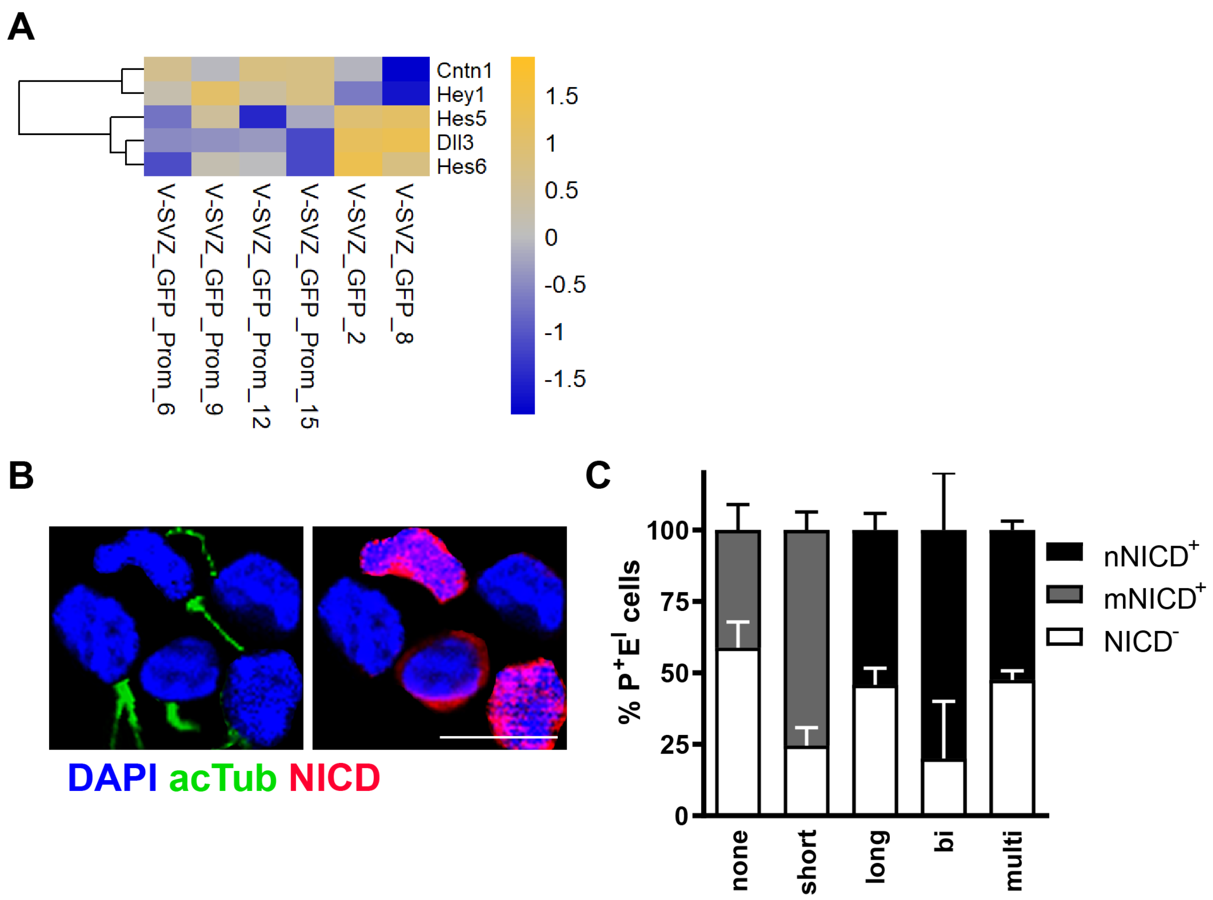


**Figure S7: Ciliated cells have activated Notch signalling.** A) Heatmap illustrating differentially regulated Notch genes. (B) Micrograph-collage of cells labelled for acetylated tubulin (acTub, green) and Notch intracellular domain (NICD, red). Scale bar indicates 10 µm. (C) Quantification of NICD-positive cells in a population of $P^{+}E^{I}$ cells, grouped according to cilia number and length. Cells are classified as not ciliated (none), short single ciliated (short), long single ciliated (long), bi-ciliated (two cilia, bi), or multi-ciliated (more than two cilia, multi). $NICD^{+}$ cells are differentiated for membrane-bound NICD (mNICD) and nuclear NICD (nNICD). Bars represent mean ± SEM. N=3.

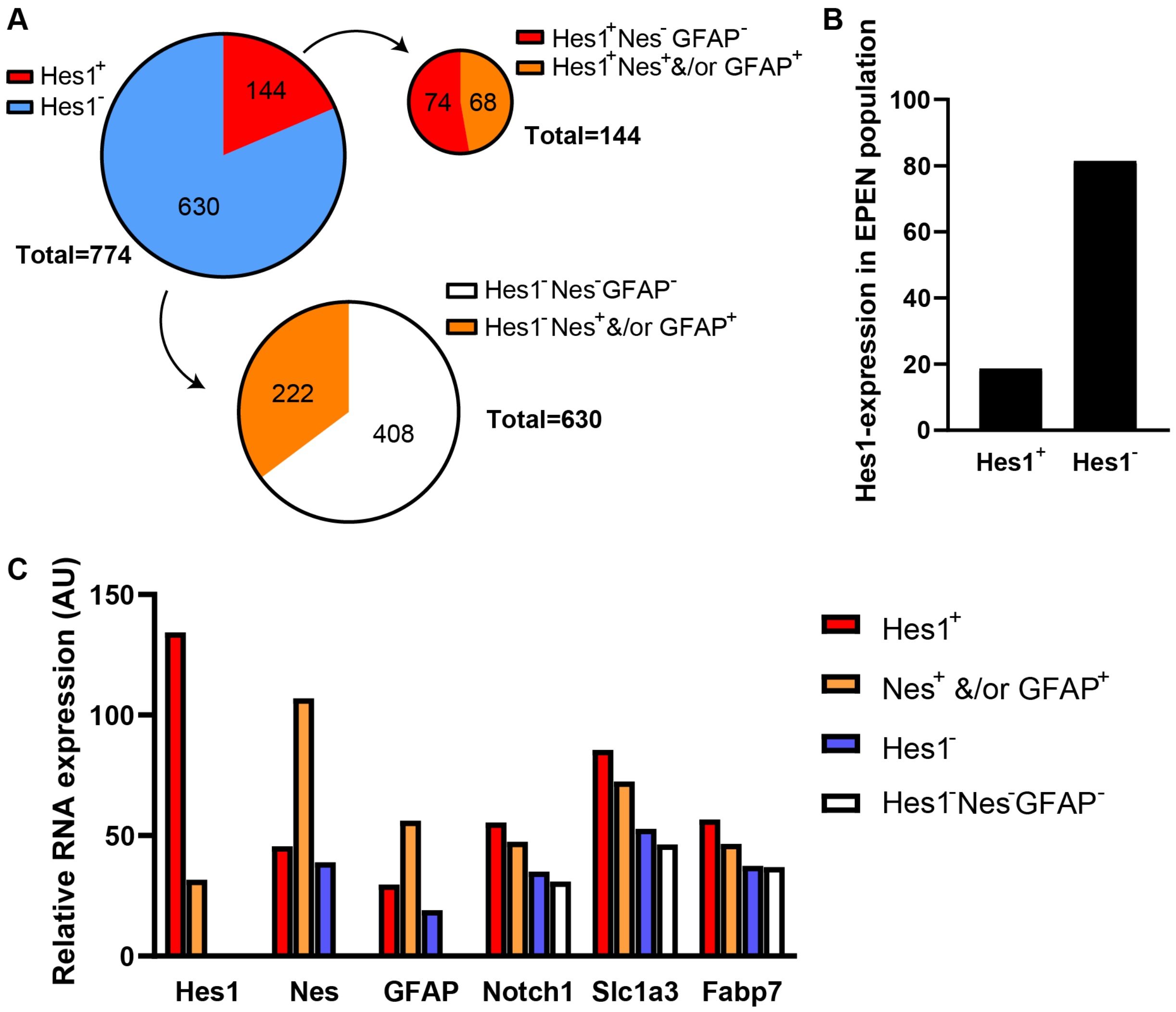


**Figure S8**

**Expression of *Hes1* is highest in a group of progenitor-like cells within the ependymal cell population.** (A) Expression of *Hes1* within cells of the ependymal cell (EPEN) population characterized on mousebrain.org, with regard to their expression of *Nestin* and/or *GFAP*. (B) Proportion of cells expressing *Hes1* in the EPEN population. (C) Relative RNA expression levels in characterized cell types shown in (A) indicates that *Hes1*-expressing cells exhibit higher expression levels of other radial glia markers like *Slc1a3* and *Fabp7* than *Hes1*-negative cells.

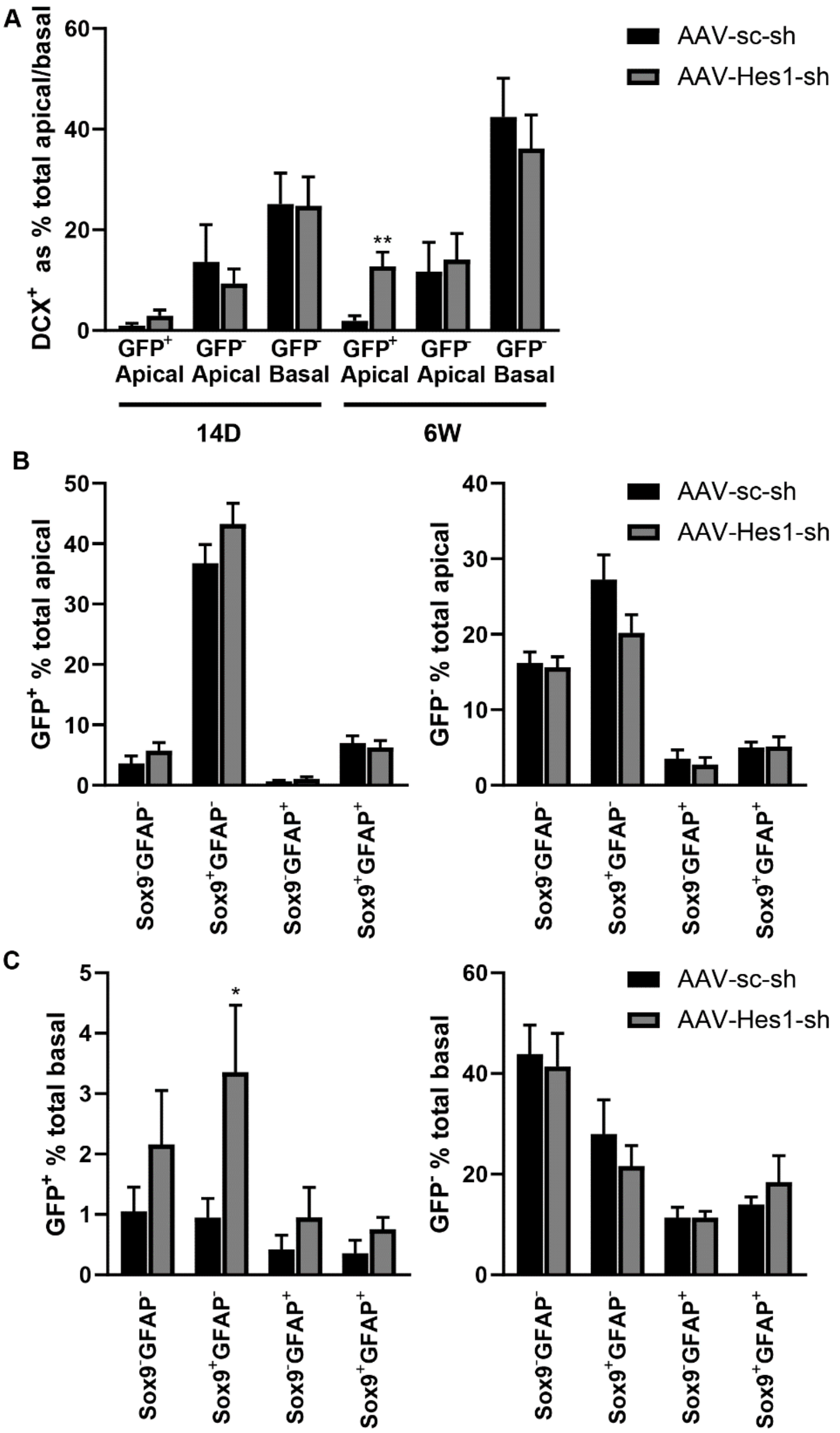


**Figure S9: Infected cells in the SVZ 14 days (14D) and 6 weeks (6W) after AAV injection.** (A) DCX-expressing cells in infected ($GFP^+$) and uninfected ($GFP^-$) apical and uninfected basal cells in the SVZ. (B+C) Expression of GFAP and Sox9 in infected and uninfected apical (B) and basal (C) cells. Bars represent mean ± SEM. * indicate significance, *$p<0.05$, **$p<0.01$. N=5 (AAV-sc-sh), N=6 (AAV-Hes1-sh).

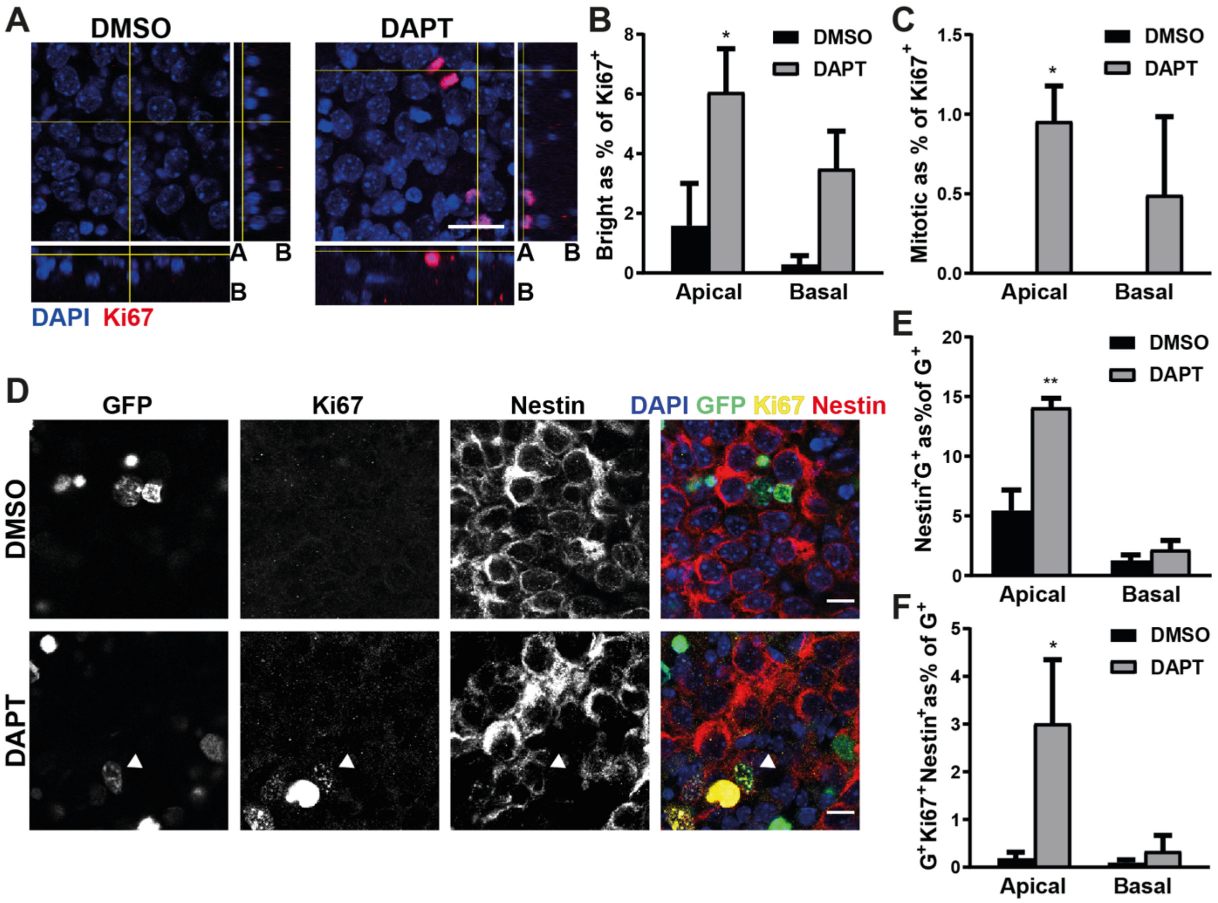


**Figure S10: Interference with the Notch signalling pathway leads to increased proliferation in apical cells.** (A) Representative micrographs of SVZ whole mounts treated with DMSO or DAPT overnight and stained with Ki67 and DAPI. Scale bar indicates 20 µm. (B,C) Quantification of apical and basal bright Ki67$^+$ cells (B) and cells undergoing mitosis (C). (D) Representative micrographs of whole mounts from hGFAP-H2BGFP mice treated with DMSO or DAPT, labelled for Ki67 and Nestin and counterstained with DAPI. Scale bars indicate 10 µm. (E,F) Quantification of cells positive for GFP, Nestin and Ki67 (E) or Nestin and GFP only (F), as percentage of total apical or basal G$^+$ cells. Bars represent mean ± SEM, * indicates significance *$p<0.05$, **$p<0.01$, ***$p<0.001$.
